# A programmable reaction-diffusion system for spatiotemporal cell signaling circuit design

**DOI:** 10.1101/2022.11.15.516470

**Authors:** Rohith Rajasekaran, Chih-Chia Chang, Elliott W. Z. Weix, Thomas M. Galateo, Scott M. Coyle

## Abstract

Cells self-organize molecules in space and time to generate complex behaviors, but we lack synthetic strategies for engineering spatiotemporal signaling. We present a programmable reaction-diffusion platform for designing protein oscillations, patterns, and circuits in mammalian cells using two bacterial proteins, MinD and MinE (MinDE). MinDE circuits act like “single-cell radios”, emitting frequency-barcoded fluorescence signals that can be spectrally isolated and analyzed using digital signal processing tools. We define how to genetically program these signals and modulate their dynamics using engineerable protein-protein interactions. By connecting MinDE to endogenous cellular pathways, we built circuits that broadcast frequency-barcoded single-cell kinase activity or that synthetically pattern actin polymerization. Our work establishes a new paradigm for probing and engineering cellular activities at length and timescales critical for biological function.

## Main Text

Cell biology is animated by the dynamic organization of protein activities in space and time. By specifying when and where specific proteins act, cells can build a diverse range of functions needed for growth, information processing, and motility from a common set of conserved molecular components (*1–3*). Disruption of this spatiotemporal organization at the single-cell level can lead to human diseases (*4–6*), and bacteria and viruses often pattern host-cell activities in new ways to hijack host-cell biology (*7–9*). Thus, the spatiotemporal organization of proteins is a key programming language of cell biology that demands deeper understanding and greater engineerable control.

Despite its fundamental importance, there are few tools that enable us to artificially organize the spatiotemporal dynamics of proteins inside living cells (*10–13*). Two common strategies are localization sequences (*14, 15*), which direct a protein of interest to a static location in the cell; and optogenetics, which allows an experimenter to organize molecules within an illuminated region of the cell (*16, 17*). Although both approaches have significant utility, they force us to choose between genetic encodability and dynamic control, failing to recapitulate the self-organization that is a hallmark of all living cells.

We sought a new paradigm: genetically encoded circuits that synthetically direct the spatiotemporal organization of proteins within a cell. To achieve this, we repurposed a positioning circuit from bacteria—the MinDE system—which is orthogonal to Eukaryotes (*18*). In *E. coli* cells, the division machinery is localized by pole-to-pole protein oscillations on the plasma membrane driven by a reaction-diffusion (*19–21*) process: nucleotide-dependent membrane association of the MinD ATPase is antagonized by its ATPase-activating protein MinE (*22, 23*). *In vitro*, MinD and MinE are sufficient to self-organize both dynamic and static protein patterns on supported lipid bilayers and can be biochemically steered towards specific patterning outcomes (*24*– *31*). In addition, minutes-timescale MinDE wave phenomena have been observed when the components are heterologously expressed in unicellular Schizosaccharomycetes fungi (*32*). These minimal requirements for patterning suggested that MinDE might produce reaction-diffusion behaviors in mammalian cells, providing a starting point for more complex spatiotemporal circuit design (Fig. 1A).

**Figure 1.**
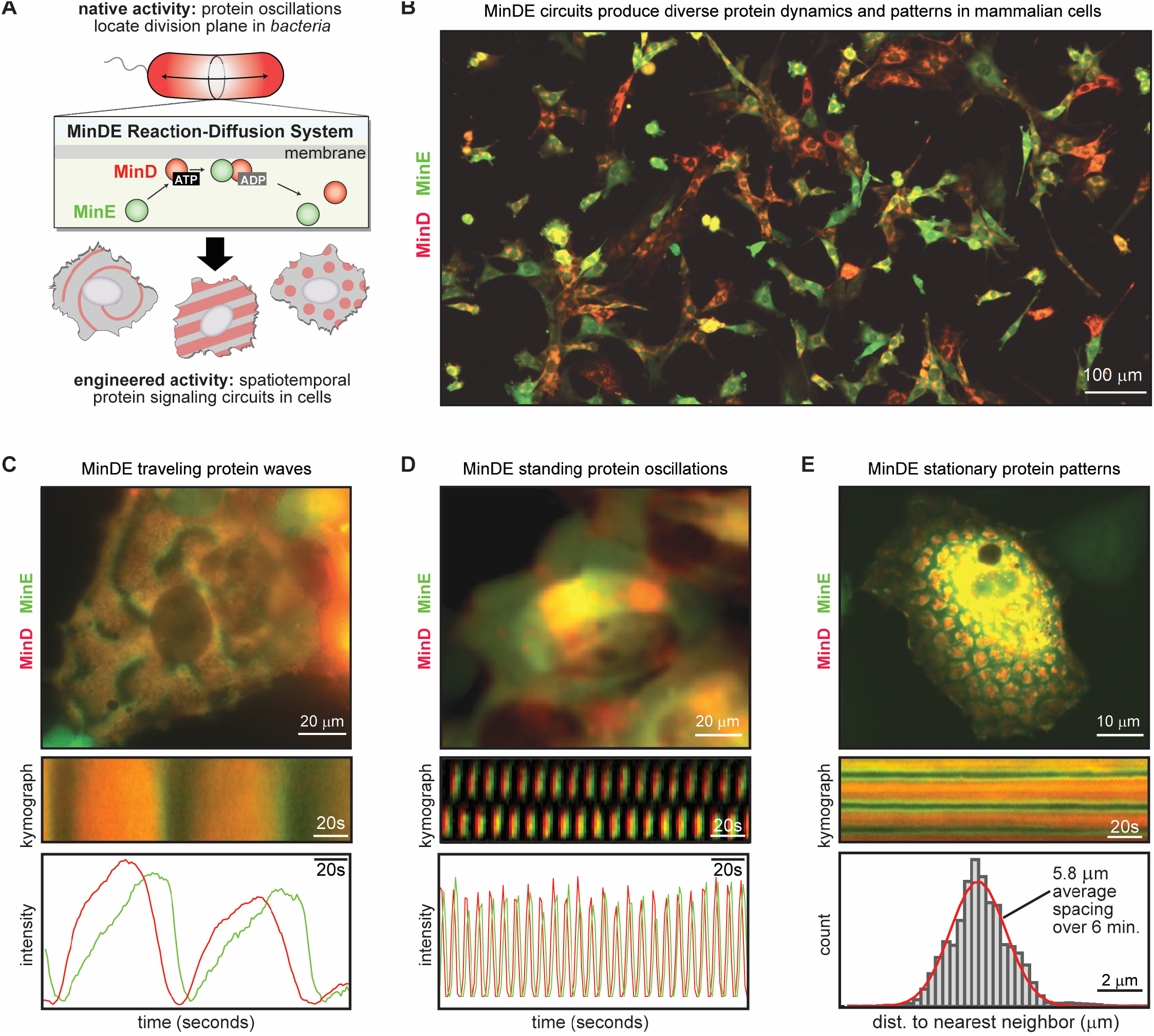
Synthetic spatiotemporal protein patterning in mammalian cells using the MinDE reaction-diffusion system. (a) The MinD ATPase and its activator MinE form a protein-based reaction-diffusion system (MinDE) in *E. coli* that generates pole-to-pole oscillations to specify the division site. Transplanting these components into human cells provides a synthetic biology platform for engineering more complex spatiotemporal signaling circuits. (b) Representative image of a population of 3T3 cells expressing mCherry-MinD and MinE-GFP at different levels. A wide array of self-organizing protein dynamics and protein patterns are seen in single cells all throughout the population. (c) Examples images, kymographs, and quantification of traveling waves (d) standing oscillations and (e) persistent stationary patterns in U2OS cells. Additional examples of MinDE reaction-diffusion phenomena across a range of different mammalian and human cell lines are detailed in Fig. S1-S2 and Movies S1-S6.

We used lentivirus to express mCherry-MinD and MinE-GFP in multiple different mammalian cell lines and imaged their spatiotemporal distribution by timelapse fluorescence microscopy. Across populations of cells harboring different MinD and MinE expression levels, we observed a stunning array of self-organizing protein dynamics and patterns inside single cells (Fig. 1B-E, Fig. S1-2, Movie S1-6). This included traveling waves, spirals, and turbulent patterns with periods ranging from seconds to minutes (Fig. 1C, Fig. S1, Movie S3); fast standing oscillations with persistent nodal structure (Fig. 1D, Fig. S1, Movie S4); and stationary Turing-type “leopard print” patterns in which regularly spaced protein domains maintained their position over time (Fig. 1E, Fig. S1, Movie S5). These MinDE patterns were distributed throughout the entire cell body, and confocal imaging suggested this whole-cell patterning was supported by MinDE interaction with the endomembrane system of the cell (Fig. S3, Movie S7). Detailed inspection of MinD and MinE fluorescence signals over time showed that for dynamic patterns, MinE slightly lagged MinD (Fig. 1B-C, Fig. S1); and for stationary patterns, domains of MinD were encircled at their perimeter by a ring of MinE (Fig. 1D, Fig. S1, Movies S2 and S5). This is the expected spatiotemporal organization of a two-component reaction-diffusion system and agrees with past *in vitro* observations of reconstituted MinDE systems (*21, 26, 30, 33, 34*).

This genetically encoded protein patterning system is powerful because its behavior can be tuned and controlled using synthetic biology and protein engineering. Thus, MinDE can provide a general platform for circuit design in which the protein patterns it generates can encode dynamic data and react to or control endogenous activities in the host cell. To analyze the content of these MinDE signals quantitatively, we took advantage of the fact that the pixel-level MinDE fluorescence dynamics at any location within a cell are strongly oscillatory (Fig. 1B-C, 2B, Fig. S1). Using Fast-Fourier Transform (FFT) (*35*), we mapped these pixel-level dynamics to the frequency-domain. The pixel-level power spectrum shows a strong peak at the frequency of the MinDE oscillation at that location (Fig. 2B). These pixel-level FFTs were used to produce an “image level power spectrum” stack, in which each slice in the image-stack corresponds to a different frequency and pixel intensity within a slice corresponds to the oscillation power of that frequency at that location (Fig. 2B). In these stacks, an individual cell is located at a particular frequency slice, indicating that a MinDE circuit generates a specific cell-wide temporal frequency. The associated phase angle of the oscillation at each pixel in that frequency slice (the phase field) provides additional information about the underlying spatial structure of that cell’s MinDE circuit, including its wavelength, defect distribution, and nodal structure (Fig. S4-5, Movie S8-9) (*36*).

**Figure 2.**
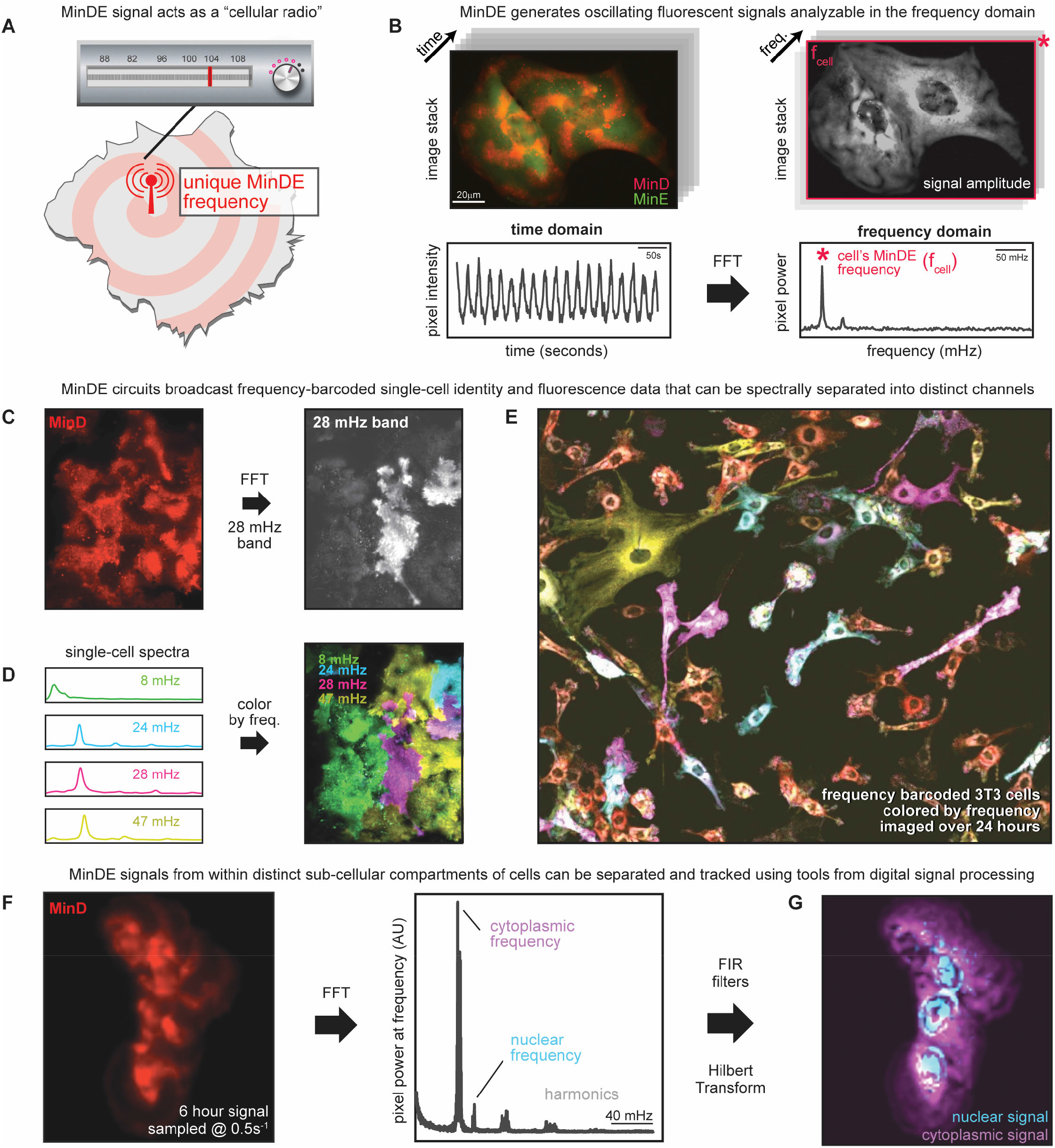
MinDE signals frequency-barcode cellular identity. (a) An oscillating MinDE signal acts like a “cellular radio station” that imparts the cell with a unique genetically encoded frequency. (b) Image-level and pixel-level time series of mCherry-MinD fluorescence data can be remapped to the frequency domain by Fast Fourier Transform (FFT) to produce image-level and pixel-level power spectra. Individual cells appear in the slice of the image-level power spectrum that corresponds to their unique MinDE cellular frequency *f*_MinDE_. (c) Frequency-domain image processing applied to the time-series data of a single fluorescent signal (mCherry-MinD). An example of spectral isolation of an individual cell with a 28 mHz MinDE frequency from its neighbors and (d) generation of a multi-channel image in which each color represents a different MinDE frequency is shown. Cells can be located based on their frequency barcode. See also Movies S10-12. (e) Representative image from a 24-hour timelapse of migrating 3T3 cells false-colored by MinDE frequency. Frequency-barcodes persist throughout the time course and enable spectral resolution of overlapping cells. See also Movie S11. (f) Separation of MinDE signals occurring in different sub-compartments of the same cell. The pixel-level power spectrum of mCherry-MinD signals collected over a 6-hour time series revealed distinct frequencies corresponding to separate nuclear and cytoplasmic MinDE oscillations. (g) Using Finite Impulse Response (FIR) filters based on (f), the nuclear and cytoplasmic signals were isolated, and the Hilbert Transform used to estimate the instantaneous power of each signal throughout the recording. This produces a multi-channel representation of the data that separately labels and tracks the cytoplasmic and nuclear signals. Identical results were obtained using Continuous Wavelet Transform. See Fig. S5 and Movies S13-17 for additional detail and examples.

Given that the temporal frequency of a cell’s MinDE signal is uniform, it provides a frequency-domain single-cell imaging barcode that acts like a “cellular radio station” (Fig. 2A). This surprising fact enabled us to perform filtering, isolation, and analysis of MinDE fluorescence signals using techniques from digital signal processing. Fluorescence from different cells within a mixed population could be isolated by simply tuning to different slices in the image power spectrum, making locating and segmenting cell boundaries trivial even at elaborate cell-cell interfaces (Fig. 2C, Fig. S2). Moreover, mixed signals arising from physically overlapping cells with distinct MinDE frequencies could be decomposed and assigned back to their cell of origin (Fig. 2D-E). For example, within a population of migrating 3T3 cells, these frequency-barcodes could label and track single cells even as they interacted and crawled on top of one another. This enabled us to produce a multi-channel rendering of the time series data in which individual cells were colored by frequency, using only a single fluorophore (Fig. 2E, Movie S10-11). This frequency-barcoding strategy was effective in all cell lines tested (Fig. S2, Movie S12). We also found that we could separate MinDE signals arising from different sub-cellular compartments within the same cell. In a 6-hour recording of a U2OS cell, the associated power spectrum showed peaks at the fundamental frequency of the MinDE oscillation in the cell body and its harmonics, as well as an additional small peak corresponding to a weak secondary MinDE oscillation in the nucleus (Fig. 2F). We separated the nuclear and cytoplasmic signals from within this single-channel recording using finite-impulse-response (FIR) filters (*35*). We then recovered the instantaneous power, phase, and frequency of the resulting signals using the Hilbert Transform (*35*), generating a multi-channel rendering that separately labels each sub-cellular compartment (Fig. 2G, Fig. S4-5, Movie S13-15). Nearly identical results were obtained using continuous wavelet transform (Fig. S5, Movie S16-17) (*37, 38*). MinDE signals thus provide an unprecedented means of encoding single-cell and sub-cellular information in synthetic protein dynamics that can be easily decoded using a suite of powerful tools from digital signal processing.

Given the utility of the signals MinDE circuits generate, we sought to program their dynamic behavior at the genetic level. We imaged thousands of MinDE circuit configurations spanning a wide range of mCherry-MinD and MinE-GFP expression levels, segmented cells based on their frequency, and extracted the average GFP and mCherry intensity within each cell as a proxy for MinE and MinD expression (Fig. 3A). Aggregating these data, we found that MinDE signal *frequency* was strongly determined by the relative levels of MinD and MinE: for a fixed MinD expression level, higher MinE levels led to higher frequencies (Fig. 3B, 3D). In contrast, the *power* (amplitude) of the MinDE signal was set by the MinD expression levels (Fig. 3C). These trends were consistent across all cell lines we analyzed (Fig. S6). Thus, a specific MinDE signal frequency and amplitude can be genetically encoded by controlling gene expression levels. These observations reflect the biochemical activities of MinD and MinE and trends seen in *in vitro:* MinD is the ATPase in the system, setting the maximum number of molecules that can participate in the oscillation; and MinE is the ATPase activator, whose concentration will determine how rapidly ATP is hydrolyzed by MinD (*18, 27, 39–42*). Consistent with this, mutant forms of MinE with faster membrane exchange (*43*) produced faster oscillations (>160 mHz) and showed different frequency-scaling relationships (Fig. 3D). This indicates that targeted mutations in MinDE components can be used to alter and engineer the mapping between gene expression levels and circuit dynamics.

**Figure 3.**
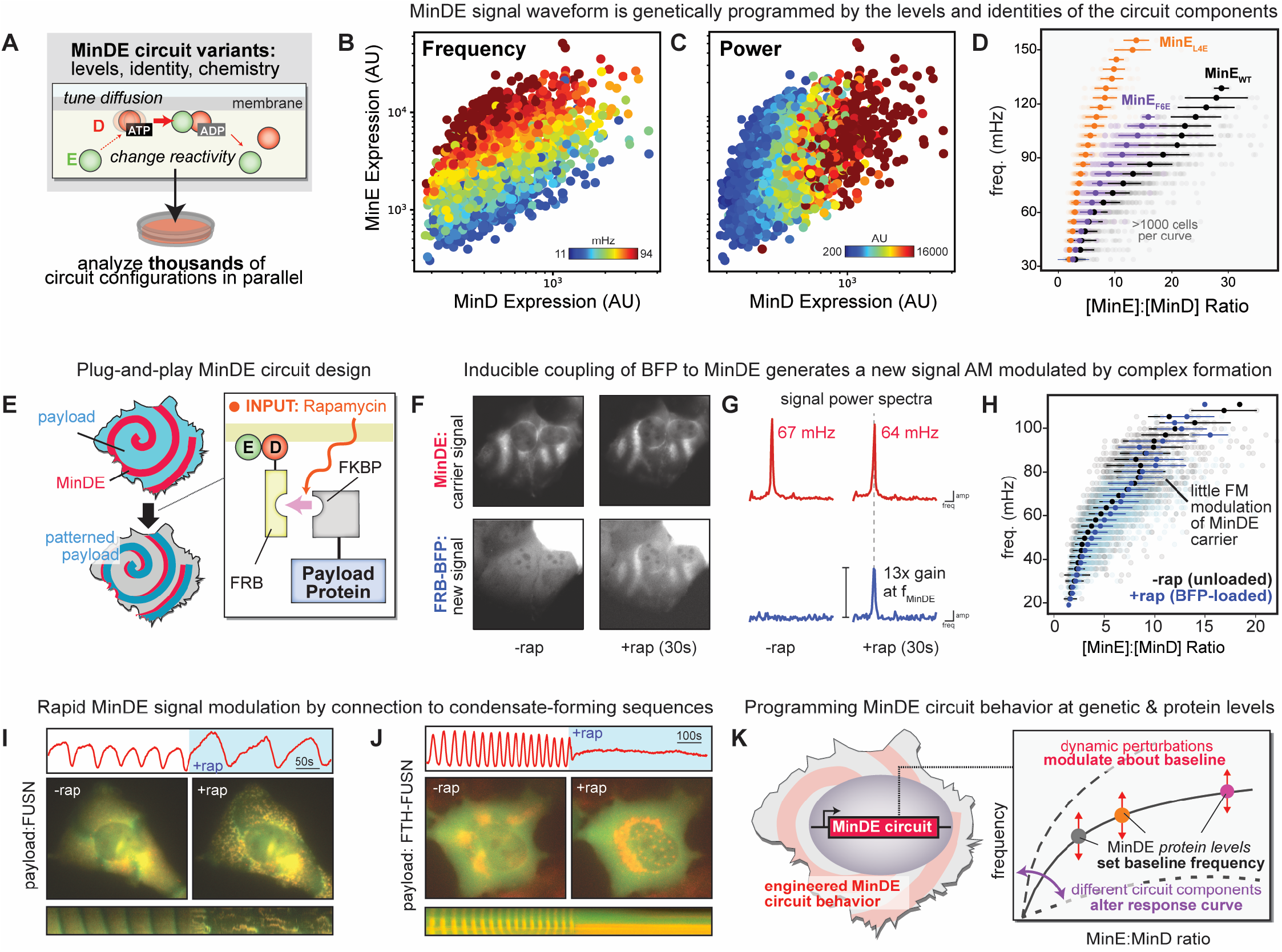
Programming and modulating MinDE circuit dynamics. (a) Thousands of MinDE circuit configurations sampling different expression levels or component ideas can be characterized in parallel using the frequency-domain image processing techniques from Fig. 2. (b) Aggregating single-cell circuit behavior across 1000s of U2OS cells reveals how MinDE circuit frequency and (c) MinDE circuit power can be programmed at a genetic level by controlling the expression levels of MinD and MinE. Similar trends were obtained across all cell-lines examined, see Fig. S6. (d) The MinE:MinD expression ratio is the major determinant of MinDE circuit frequency. This enables characterization and comparison of the frequency-scaling behavior of MinDE circuits harboring different mutant components. The frequency-scaling behavior for MinE mutants that affect MinE membrane affinity and diffusivity are shown. See also Fig. S6. (e) Schematic of a plug-and-play platform for inducible recruitment of protein payloads to MinDE circuits based on rapamycin-dependent interaction between FRB-MinD and FKBP-payloads. (f) Inducible recruitment of a simple BFP payload using the system described in (e). Representative FRB-mCherry-MinD and BFP-FKBP images of cells pre and post rapamycin inductions are shown, demonstrating rapamycin-dependent colocalization of BFP with MinD. See also Movie S18. (g) Power-spectra for the MinD and BFP signals in (f) pre and post rapamycin induction. MinD shows a consistent high-power signal in the presence or absence of rapamycin. The BFP signal has little power at the MinD frequency in the absence of rapamycin, but high-power (13x increase) when rapamycin is added. (h) MinDE frequency scaling behavior for the MinD-BFP recruitment circuit in the presence or absence of rapamycin. There is little change in the steady-state frequency of the MinD signal upon recruitment. (i) Representative images, pixel-level intensity time course, and kymograph pre and post recruitment of the condensate-forming intrinsically disordered region of FUSN to MinDE. The MinDE signal undergoes rapid frequency modulation as puncta-like structures assemble and disassemble along the wave trajectory. Additional examples and data in Fig. S7 and Movie S19. (j) As in (i) but using a pre-formed protein condensate (FTH-FUSN) as a payload. MinDE oscillations immediately arrest as MinDE components colocalize into droplets. Additional examples and data in Fig. S7 and Movie S19. (k) Schematic for how MinDE circuit behaviors and dynamics can be genetically programmed and modulated by engineered protein-protein interactions.

Because MinDE circuits are fast and protein-based, more complex signaling dynamics or composite signals can be created by coupling MinDE to other proteins or cellular structures. We created a plug-and-play platform that dynamically connects MinDE to other components, based on the chemically inducible heterodimerization system FKBP/FRB (Fig. 3E) (*44*). When FRB was fused to MinD, an FKBP-BFP test payload did not colocalize with MinDE in the absence of rapamycin. However, within seconds of adding rapamycin, the BFP signal colocalized with MinDE in space and time (Fig. 3F, Movie S18). As a result, this circuit generates a new oscillatory signal in the BFP channel whose frequency matches the MinDE carrier signal but whose amplitude is modulated in real time by the FKBP/FRB interaction (Fig. 3G).

As MinDE interacts with other proteins in the cell, the spatiotemporal dynamics of the MinDE signal itself can also change in response. For a simple BFP payload, we observed little change in the steady-state frequency of the MinDE signal upon loading (Fig. 3H); that is, the derived BFP signal largely inherits the frequency of the MinDE carrier it arises from. We then tested the effects of connecting MinDE to well-characterized “intrinsically disordered region” sequences (IDRs) that have different propensities to form liquid condensates, gels, or aggregates with altered diffusivity (*45-47*). While weak IDRs like RGG had little effect on MinDE dynamics, recruitment of stronger IDRs derived from FUSN or DDX4 rapidly altered MinDE spatiotemporal behavior (Fig. 3I, Fig. S7, Movies S19). For these circuits, oscillating puncta assembly and disassembly were observed along the wave trajectory, coincident with a drastic decrease in the circuit frequency and redistribution of circuit power. In contrast, connection of MinDE circuits to large, preformed protein-condensates (via FTH1-FUSN (*47*)) or to the microtubule cytoskeleton (via TPPP (*48*)) led to immediate arrest of MinDE oscillations (Fig. 3J, Fig. S7, Movie S19).

Taken together, these results indicate that the baseline MinDE spatiotemporal dynamics can be programmed at a genetic level and further modulated in real-time at the protein level through dynamic connection to other cellular components (Fig. 3K). Depending on the nature of these interactions, MinDE dynamics can persist and drive other processes (such as condensate assembly and disassembly) or be overridden by stronger localization signals. Thus, the space of possibilities for sculpting MinDE behavior in living cells is vast, creating a new frontier for synthetic biology to experimentally explore reaction-diffusion dynamics, patterning, and signal generation.

We next applied this knowledge to design specific MinDE circuits for understanding or engineering cell biology. Taking advantage of the “radio”-like properties of MinDE signals, we first created circuits that read out and broadcast frequency-barcoded dynamic cell-state data, using protein kinase A (PKA) signaling activity as a test case (Fig. 4A). For this circuit, we fused a substrate sequence for PKA to MinD and co-expressed this with mCerulean-FHA, which specifically binds to the phosphorylated substrate sequence (*49*). This exact substrate/reader pair was previously used to develop a FRET-based PKA activity reporter (*50*). The resulting MinDE circuit generates a new signal derived from MinDE in the mCerulean channel whose amplitude is dynamically modulated by PKA signaling activity. In resting cells, this PKA activity signal had low power as there was little colocalization of FHA-mCerulean with MinDE (Fig. 4B, Fig. S8). However, upon stimulation with the PKA agonist isoprenaline, FHA immediately began co-oscillating with the MinDE carrier, leading to increase in the power of the PKA activity signal at the pixel level (Fig. 4B, Fig. S8). Importantly, the ratio of power between the MinDE carrier signal and the PKA activity signal was consistent everywhere within the cell, facilitating normalization, comparison and aggregation of the single-pixel signaling trajectories (Fig. 4B, Fig. S8, Movie S20-21). For a saturating dose of isoprenaline, the increase in this power ratio could be as much as 600%, more than twenty-fold the reported dynamic range of the FRET sensor from which the circuit components were derived (*50*).

**Figure 4.**
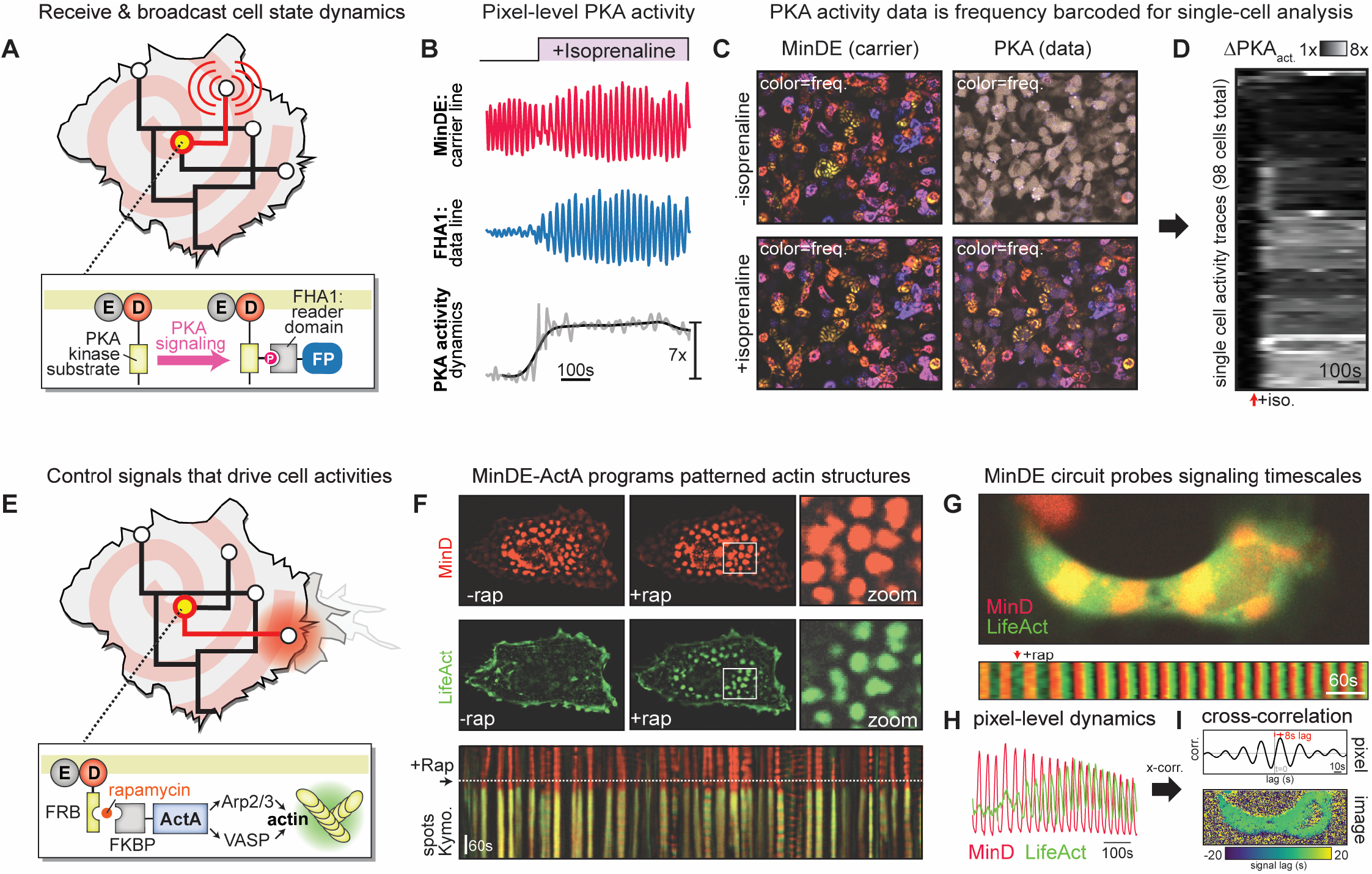
Engineering MinDE circuits to broadcast or control dynamic cellular activities. (a) MinDE circuits can be engineered to receive, barcode, and broadcast the dynamics of any fluorescent cell-state reporter. A specific circuit that broadcasts PKA kinase signaling activity was designed by fusing a PKA substrate to MinD and co-expressing FHA1-mCerulean, which binds the phosphorylated PKA substrate. This PKA→MinDE→FHA1 circuit creates a new signal derived from MinDE (the carrier) in the mCerulean channel that is modulated by PKA signaling activity (the data line). (b) Example of pixel-level read-out of PKA signaling activity in a 293T cell using the circuit in (a). Upon treatment with the PKA agonist isoprenaline, the FHA1 signal (PKA data line) begins to co-oscillate with MinDE (carrier signal). Normalization of the power of the FHA1 signal to the MinDE carrier recovers the real-time PKA signaling dynamics for that pixel within the cell. (c) Representative images of a population of cells bearing the circuit in (a), showing the power of the MinD (carrier line) or FHA1 (PKA data line) signals pre or post treatment with isoprenaline, color-coded by frequency. (d) 98 single-cell PKA signaling trajectories extracted using the frequency-barcoded data broadcast by the cells in (c), clustered by response behavior. Additional detail, data, and examples in Fig. S8 and Movies S20-S21. (e) MinDE circuits can also be engineered to act as control signals that drive cellular activities. A specific circuit that inducibly couples MinDE spatiotemporal dynamics to actin polymerization upon rapamycin induction was designed using the bacterial signaling effector ActA as an FKBP-payload. (f) Demonstration that the MinDE→ActA control signal circuit can organize the polymerization of actin into structures with position and shape dictated by the MinDE signal, using a stationary MinDE pattern as an example control signal. Representative images for MinD (control signal) and the LifeAct actin reporter (output signal), pre (-rap) and post (+rap) coupling are shown, along with a kymograph passing through multiple MinDE spots. (g) When MinDE control signals are dynamic, they can be used to probe signaling timescales within the cell. An example cell bearing a MinDE→ActA circuit driven by a dynamic MinD wave is shown along with an associated kymograph and (h) a single-pixel intensity time course of the MinD and LifeAct signals indicate that the LifeAct (output signal). (i) Representative pixel-level cross-correlation profile between MinD and LifeAct signals and associated image-level rendering of the extracted time-lag across the field of view in (g). This shows a consistent 8s lag for signal transmission from the MinDE-ActA circuit to actin everywhere within the cell. Additional examples and analyses are detailed in Fig. S9-11 and Movie S22-23.

Critically, this readout of PKA signaling is barcoded by the cell’s underlying MinDE frequency (Fig. 4C), allowing the PKA activity for individual cells to be spectrally isolated from others in the frequency domain. This enabled rapid, high-throughput extraction of single-cell PKA signaling trajectories from fields of cells even at low magnification (Fig. 4D). This ability to couple cellular data to the MinDE frequency-barcode is not possible with other imaging barcoding strategies based on ratioing multiple fluorescent proteins (*51, 52*). Moreover, because only on a single fluorophore is needed to track the MinDE carrier signal, it is possible to barcode and broadcast multiple data lines for different signaling pathways or activity reporters in parallel using this approach. To our knowledge, there are no existing strategies that provide the barcoding, amplification, and multiplexing potential of MinDE cellular radio circuits for reading out single-cell data in real time.

We then explored whether the same MinDE signals we used to broadcast endogenous cellular activities could also be used to synthetically reorganize them. We built circuits that targeted a pathway that would produce an output signal clearly localized in space and time: actin polymerization. Many endogenous signaling pathways and bacterial effector proteins organize actin polymerization in the cell through signaling to the Arp2/3 complex (*6, 53, 54*). A fragment of one such effector from *Listeria*, ActA, can support synthetic actin polymerization when clustered on surfaces or membranes (*9, 55*). Using our plug-and-play inducible system, we generated cell lines in which an ActA payload could be coupled to MinDE spatiotemporal patterns and used LifeAct-GFP (*56*) to monitor the associated actin dynamics in real-time (Fig. 4E). This allowed us to query a range of MinDE-to-ActA control signals for their ability to support and organize actin polymerization. In cells displaying stationary MinDE patterns of geometric spots, the LifeAct-GFP signal did not colocalize with MinDE patterns in the absence of rapamycin. However, upon addition of rapamycin, we observed rapid production of filamentous actin structures with size, shape and position controlled by the associated MinDE pattern (Fig 4F, Fig. S9-10, Movie S22-23). This clearly shows that the spatiotemporal organization encoded by a MinDE circuit can be used as a blueprint to productively localize host-cell signaling activities in space and time.

Interestingly, when ActA was connected to *dynamic* MinDE patterns, LifeAct-GFP oscillated at the same frequency as MinDE but with a noticeable delay between the two signals (Fig. 4G-H, Fig. S11). Cross-correlation (*35* of the oscillatory LifeAct and MinDE signals defined a time-lag of 6-10 seconds in the cells we inspected (Fig. 4I, Fig. S11, Movie S23), which we interpret as the time it takes for MinDE recruitment of ActA to drive productive signaling to the actin polymerization machinery in our circuit context. We emphasize that this specific timescale likely depends on the expression levels of ActA, the power of the MinDE oscillation, and the availability of actin regulatory components in any individual cell. Nevertheless, it demonstrates that MinDE signals not only can act as genetically encoded spatiotemporal controllers of the cell, but can also dynamically probe spatiotemporal signaling constraints on the pathways that we connect them to.

We have shown that a bacterial reaction-diffusion system, MinDE, can be used as an orthogonal synthetic biology platform to generate self-organizing, genetically encoded protein oscillations, patterns, and spatiotemporal signaling circuits in mammalian cells. The oscillatory nature of MinDE signals enables them to be deployed as cellular radios that barcode single-cell identity or compartments with a unique frequency. Capitalizing on these properties, we developed specific circuits that can read out and control mammalian cell biology with subcellular resolution. Our work provides a foundation for building more complex protein-based reaction-diffusion circuits that incorporate feedback and sense data into their spatiotemporal dynamics, providing a new suite of tools for visualizing, probing, and perturbing cell biology. It also provides an experimental test-bed for leveraging synthetic biology to experimentally explore the physics and chemistry of reaction-diffusion systems that incorporate time-varying protein-levels, localization, or diffusivity (*57*). Taken together, we establish a new paradigm for synthetic interaction with cells at length and timescales critical to biological function.

## Acknowledgements

We thank members of the Coyle Lab, A. Weeks, W. Bement, A. Larson, L. Lui, L. Pack, and K. Roybal for advice, helpful discussions, and critical reading of the manuscript.

## Author contributions

RR and SMC conceived the overall project. CCC and SMC conceived the experiments involving condensate-forming sequences. The experimental plan was implemented by RR, CCC, EWZW, TMG, and SMC. RR collected all the data in the paper except for that in Fig. 3I-J, Fig. S7, and Movie S18, which were performed by CCC and EWZW. TMG assisted RR in the collection and analysis of data for Fig. 3D. RR and SMC developed the frequency-domain image analysis pipelines and code. RR and SMC prepared the figures and wrote the manuscript with suggestions from all authors. SMC supervised all aspects of the work.

## Data and materials availability

All data and analysis code are available in the manuscript, supplementary materials, and the Coyle-Lab GitHub repository. Reagents are available from the corresponding author upon reasonable request. Plasmids from this paper will be made available on Addgene. A playlist containing all supplementary movies in a streaming format can be found at: http://youtu.be/watch?v=KDgHVrjdydY&list=PLx2v1NUlEZF_idH1goDHYimtlP8c7mnLB&index=1

## References

1. J. E. Purvis, G. Lahav, Encoding and Decoding Cellular Information through Signaling Dynamics. Cell. 152, 945–956 (2013).

2. T. D. Pollard, G. G. Borisy, Cellular Motility Driven by Assembly and Disassembly of Actin Filaments. Cell. 112, 453–465 (2003).

3. E. Karsenti, Self-organization in cell biology: a brief history. Nat Rev Mol Cell Bio. 9, 255–262 (2008).

4. L. J. Bugaj, A. J. Sabnis, A. Mitchell, J. E. Garbarino, J. E. Toettcher, T. G. Bivona, W. A. Lim, Cancer mutations and targeted drugs can disrupt dynamic signal encoding by the Ras-Erk pathway. Science. 361 (2018), doi:10.1126/science.aao3048.

5. D. Yamazaki, S. Kurisu, T. Takenawa, Regulation of cancer cell motility through actin reorganization. Cancer Sci. 96, 379–386 (2005).

6. M. Symons, J. M. J. Derry, B. Karlak, S. Jiang, V. Lemahieu, F. McCormick, U. Francke, A. Abo, Wiskott–Aldrich Syndrome Protein, a Novel Effector for the GTPase CDC42Hs, Is Implicated in Actin Polymerization. Cell. 84, 723–734 (1996).

7. J. E. Galán, Common Themes in the Design and Function of Bacterial Effectors. Cell Host Microbe. 5, 571–579 (2009).

8. A. Sodhi, S. Montaner, J. S. Gutkind, Viral hijacking of G-protein-coupled-receptor signalling networks. Nat Rev Mol Cell Bio. 5, 998– 1012 (2004).

9. L. A. Cameron, M. J. Footer, A. van Oudenaarden, J. A. Theriot, Motility of ActA protein-coated microspheres driven by actin polymerization. Proc National Acad Sci. 96, 4908–4913 (1999).

10. A. H. Chau, J. M. Walter, J. Gerardin, C. Tang, W. A. Lim, Designing Synthetic Regulatory Networks Capable of Self-Organizing Cell Polarization. Cell. 151, 320–332 (2012).

11. M. B. Elowitz, S. Leibler, A synthetic oscillatory network of transcriptional regulators. Nature. 403, 335–338 (2000).

12. B. Jayanthi, B. Bachhav, Z. Wan, S. M. Legaspi, L. Segatori, A platform for post-translational spatiotemporal control of cellular proteins. Synthetic Biology. 6, ysab002 (2021).

13. S. Regot, J. J. Hughey, B. T. Bajar, S. Carrasco, M. W. Covert, High-Sensitivity Measurements of Multiple Kinase Activities in Live Single Cells. Cell. 157, 1724–1734 (2014).

14. J. F. Hancock, K. Cadwallader, H. Paterson, C. J. Marshall, A CAAX or a CAAL motif and a second signal are sufficient for plasma membrane targeting of ras proteins. Embo J. 10, 4033–4039 (1991).

15. A. Lange, R. E. Mills, C. J. Lange, M. Stewart, S. E. Devine, A. H. Corbett, Classical Nuclear Localization Signals: Definition, Function, and Interaction with Importin α. J Biol Chem. 282, 5101–5105 (2007).

16. J. E. Toettcher, O. D. Weiner, W. A. Lim, Using Optogenetics to Interrogate the Dynamic Control of Signal Transmission by the Ras/Erk Module. Cell. 155, 1422–1434 (2013).

17. A. Levskaya, O. D. Weiner, W. A. Lim, C. A. Voigt, Spatiotemporal control of cell signalling using a light-switchable protein interaction. Nature. 461, 997–1001 (2009).

18. J. Lutkenhaus, The ParA/MinD family puts things in their place. Trends Microbiol. 20, 411–418 (2012).

19. A. M. Turing, The chemical basis of morphogenesis. Philos. Trans. R. Soc. Lond., B, Biol. Sci. 237, 37–72 (1952).

20. S. Kondo, T. Miura, Reaction-Diffusion Model as a Framework for Understanding Biological Pattern Formation. Science. 329, 1616– 1620 (2010).

21. B. Ramm, T. Heermann, P. Schwille, The E. coli MinCDE system in the regulation of protein patterns and gradients. Cell Mol Life Sci. 76, 4245–4273 (2019).

22. D. M. Raskin, P. A. J. de Boer, MinDE-Dependent Pole-to-Pole Oscillation of Division Inhibitor MinC in Escherichia coli. J Bacteriol. 181, 6419–6424 (1999).

23. K. C. Huang, Y. Meir, N. S. Wingreen, Dynamic structures in Escherichia coli: Spontaneous formation of MinE rings and MinD polar zones. Proc National Acad Sci. 100, 12724–12728 (2003).

24. M. Loose, E. Fischer-Friedrich, J. Ries, K. Kruse, P. Schwille, Spatial Regulators for Bacterial Cell Division Self-Organize into Surface Waves in Vitro. Science. 320, 789–792 (2008).

25. F. Brauns, G. Pawlik, J. Halatek, J. Kerssemakers, E. Frey, C. Dekker, Bulk-surface coupling identifies the mechanistic connection between Min-protein patterns in vivo and in vitro. Nat Commun. 12, 3312 (2021).

26. P. Glock, B. Ramm, T. Heermann, S. Kretschmer, J. Schweizer, J. Mücksch, G. Alagöz, P. Schwille, Stationary Patterns in a Two-Protein Reaction-Diffusion System. Acs Synth Biol. 8, 148–157 (2019).

27. S. Kretschmer, L. Harrington, P. Schwille, Reverse and forward engineering of protein pattern formation. Philosophical Transactions Royal Soc B Biological Sci. 373, 20170104 (2018).

28. S. Kretschmer, T. Heermann, A. Tassinari, P. Glock, P. Schwille, Increasing MinD’s Membrane Affinity Yields Standing Wave Oscillations and Functional Gradients on Flat Membranes. Acs Synth Biol. 10, 939–949 (2021).

29. P. Glock, F. Brauns, J. Halatek, E. Frey, P. Schwille, Design of biochemical pattern forming systems from minimal motifs. Elife. 8, e48646 (2019).

30. A. G. Vecchiarelli, M. Li, M. Mizuuchi, L. C. Hwang, Y. Seol, K. C. Neuman, K. Mizuuchi, Membrane-bound MinDE complex acts as a toggle switch that drives Min oscillation coupled to cytoplasmic depletion of MinD. Proc National Acad Sci. 113, E1479–E1488 (2016).

31. K. Mizuuchi, A. G. Vecchiarelli, Mechanistic insights of the Min oscillator via cell-free reconstitution and imaging. Phys Biol. 15, 031001 (2018).

32. B. Ramm, A. Goychuk, A. Khmelinskaia, P. Blumhardt, H. Eto, K. A. Ganzinger, E. Frey, P. Schwille, A diffusiophoretic mechanism for ATP-driven transport without motor proteins. Nat Phys. 17, 850–858 (2021).

33. V. Ivanov, K. Mizuuchi, Multiple modes of interconverting dynamic pattern formation by bacterial cell division proteins. Proc National Acad Sci. 107, 8071–8078 (2010).

34. M. Loose, E. Fischer-Friedrich, J. Ries, K. Kruse, P. Schwille, Spatial Regulators for Bacterial Cell Division Self-Organize into Surface Waves in Vitro. Science. 320, 789–792 (2008).

35. L. R. R. and B. Gold, Theory and Application of Digital Signal Processing (1975).

36. T. H. Tan, J. Liu, P. W. Miller, M. Tekant, J. Dunkel, N. Fakhri, Topological turbulence in the membrane of a living cell. Nat Phys. 16, 657–662 (2020).

37. S. Sinha, P. S. Routh, P. D. Anno, J. P. Castagna, Spectral decomposition of seismic data with continuous-wavelet transform. Geophysics. 70, P19–P25 (2005).

38. G. Thakur, E. Brevdo, N. S. Fučkar, H.-T. Wu, The Synchrosqueezing algorithm for time-varying spectral analysis: Robustness properties and new paleoclimate applications. Signal Process. 93, 1079–1094 (2013).

39. H. Zhou, R. Schulze, S. Cox, C. Saez, Z. Hu, J. Lutkenhaus, Analysis of MinD Mutations Reveals Residues Required for MinE Stimulation of the MinD ATPase and Residues Required for MinC Interaction. J Bacteriol. 187, 629–638 (2005).

40. K. Park, W. Wu, S. Lovell, J. Lutkenhaus, Mechanism of the asymmetric activation of the MinD ATPase by MinE. Mol Microbiol. 85, 271–281 (2012).

41. V. Ivanov, K. Mizuuchi, Multiple modes of interconverting dynamic pattern formation by bacterial cell division proteins. Proc National Acad Sci. 107, 8071–8078 (2010).

42. M. Bonny, E. Fischer-Friedrich, M. Loose, P. Schwille, K. Kruse, Membrane Binding of MinE Allows for a Comprehensive Description of Min-Protein Pattern Formation. Plos Comput Biol. 9, e1003347 (2013).

43. S. Kretschmer, K. Zieske, P. Schwille, Large-scale modulation of reconstituted Min protein patterns and gradients by defined mutations in MinE’s membrane targeting sequence. Plos One. 12, e0179582 (2017).

44. J. Choi, J. Chen, S. L. Schreiber, J. Clardy, Structure of the FKBP12-Rapamycin Complex Interacting with Binding Domain of Human FRAP. Science. 273, 239–242 (1996).

45. S. F. Banani, H. O. Lee, A. A. Hyman, M. K. Rosen, Biomolecular condensates: organizers of cellular biochemistry. Nat Rev Mol Cell Bio. 18, 285–298 (2017).

46. Y. Shin, J. Berry, N. Pannucci, M. P. Haataja, J. E. Toettcher, C. P. Brangwynne, Spatiotemporal Control of Intracellular Phase Transitions Using Light-Activated optoDroplets. Cell. 168, 159-171.e14 (2017).

47. D. Bracha, M. T. Walls, M.-T. Wei, L. Zhu, M. Kurian, J. L. Avalos, J. E. Toettcher, C. P. Brangwynne, Mapping Local and Global Liquid Phase Behavior in Living Cells Using Photo-Oligomerizable Seeds. Cell. 175, 1467-1480.e13 (2018).

48. M. Fu, T. S. McAlear, H. Nguyen, J. A. Oses-Prieto, A. Valenzuela, R. D. Shi, J. J. Perrino, T.-T. Huang, A. L. Burlingame, S. Bechstedt, B. A. Barres, The Golgi Outpost Protein TPPP Nucleates Microtubules and Is Critical for Myelination. Cell. 179, 132-146.e14 (2019).

49. D. Durocher, I. A. Taylor, D. Sarbassova, L. F. Haire, S. L. Westcott, S. P. Jackson, S. J. Smerdon, M. B. Yaffe, The Molecular Basis of FHA Domain:Phosphopeptide Binding Specificity and Implications for Phospho-Dependent Signaling Mechanisms. Mol Cell. 6, 1169–1182 (2000).

50. J. Zhang, C. J. Hupfeld, S. S. Taylor, J. M. Olefsky, R. Y. Tsien, Insulin disrupts β-adrenergic signalling to protein kinase A in adipocytes. Nature. 437, 569–573 (2005).

51. J. Livet, T. A. Weissman, H. Kang, R. W. Draft, J. Lu, R. A. Bennis, J. R. Sanes, J. W. Lichtman, Transgenic strategies for combinatorial expression of fluorescent proteins in the nervous system. Nature. 450, 56–62 (2007).

52. B. Richier, I. Salecker, Versatile genetic paintbrushes: Brainbow technologies. Wiley Interdiscip Rev Dev Biology. 4, 161–180 (2015).

53. E. D. Goley, M. D. Welch, The ARP2/3 complex: an actin nucleator comes of age. Nat Rev Mol Cell Bio. 7, 713–726 (2006).

54. M. D. Welch, J. Rosenblatt, J. Skoble, D. A. Portnoy, T. J. Mitchison, Interaction of Human Arp2/3 Complex and the Listeria monocytogenes ActA Protein in Actin Filament Nucleation. Science. 281, 105–108 (1998).

55. H. Nakamura, E. Rho, D. Deng, S. Razavi, H. T. Matsubayashi, T. Inoue, Biorxiv, in press, doi:10.1101/2020.03.30.016360.

56. J. Riedl, A. H. Crevenna, K. Kessenbrock, J. H. Yu, D. Neukirchen, M. Bista, F. Bradke, D. Jenne, T. A. Holak, Z. Werb, M. Sixt, R. Wedlich-Soldner, Lifeact: a versatile marker to visualize F-actin. Nat Methods. 5, 605–607 (2008).

57. J. Halatek, E. Frey, Rethinking pattern formation in reaction–diffusion systems. Nat Phys. 14, 507–514 (2018).

