## Supplemental Materials and Methods for "A programmable reaction-diffusion system for spatiotemporal cell signaling circuit design"

**This PDF file contains:**

Materials and Methods.
Supplemental Figures S1-S11
Supplemental Movie Legends

### Materials and Methods

Gene synthesis and cloning

Genes encoding *E. coli* MinD with N-terminal mCherry fusion and MinE with C-terminal EGFP fusion were obtained by gene synthesis as clonal genes (Twist Bioscience). Gene fragments were codon optimized for expression in human cells using Benchling's website tool. The genes were subsequently subcloned into pLV-EF1a-IRES lentiviral transfer plasmids and sequences were verified before use. MinE single-point mutants were generated by standard molecular biology methods.

Gene encoding N-terminal FRB fusion to mCherry-MinD was generated by Gibson homology cloning. Fragment encoding FRB was generated via PCR using an existing template (addgene #31181). Gene encoding BFP-FKBP was obtained as gblocks (Integrated DNA Technologies).

Human FTH1 (Ferritin Heavy Chain-1), TPPP (Tubulin Polymerization Promoting Protein), DDX4 (residues 1-236), and LAF-1 RGG (residues 1-200) fragments were obtained as gBlocks (Integrated DNA Technologies). Human FUS (residues 1-214) was amplified by PCR using pHR-FUSN-mCh-Cry2WT (addgene #101223).

Fragment encoding the substrate for PKA (LRRATLVD) was obtained as single stranded DNA oligomer (Integrated DNA Technologies) and N-terminal fusion to mCherry-MinD was generated by standard molecular biology techniques. Cognate binding partner to the phosphorylated PKA substrate was obtained from addgene (#138202).

Actuator construct composed of *Listeria* ActA(residues 1-584)-BFP-FKBP was obtained by gene synthesis as clonal gene (Twist Bioscience) in a SFFV-promoter lentiviral transfer plasmid (doi: 10.1101/2020.03.30.016360). Gene fragments were codon optimized for expression in human cells using Benchling's website tool. LifeAct-GFP reporter was obtained from addgene (#51010).

Lentiviral transduction of mammalian cell lines

Pantropic VSV-G pseudotyped lentivirus was produced by transfecting 293T cells (ATCC CRL-3216) with a pLV-EF1a-IRES or pTwist-SFFV transgene expression vector and the viral packaging plasmids psPAX2 and pMD2.G using Fugene HD (Promega #E2312). Viral production was performed in 6-well tissue culture treated plates (Corning 3335). After 72 hours, viral supernatant was harvested, filtered using 0.45μM PES syringe filter, and added to mammalian cell lines with 2μg/ml Polybrene transfection reagent (Sigma TR-1003-G). When necessary, viral supernatant was concentrated by centrifugal filtration in a 100k MWCO PES Spin-X UF 20 Concentrator (Corning 431491) at 3214xg. Mammalian cells were exposed to the virus for 24 hr. After 5 days, cells were sorted on an FACSAria cell sorter (BD Biosciences) based on fluorescent protein expression levels. Cells were sorted to achieve diverse expression levels. Mammalian cells were expanded for 7 days before use in microscopy experiments. When necessary, cells were expanded for 24 hours after removal of viral particles and imaged immediately.

Cancer cell lines culturing protocols

Mammalian cells were cultured to confluence in Dulbecco's Modified Eagle's Medium (DMEM) - high glucose (Sigma D6429) supplemented with 10% FBS (Cytiva Life Sciences SH30396.03) and 1% Penicillin-Streptomycin (ThermoFisher 15140122). At each passage, adherent cells were washed with PBS (at 37 °C) and TrypLE (ThermoFisher Scientific 12604021) was added to detach the cells from the flask surface. Flasks were incubated at 37 °C until the cells detached, typically 5 to 10 min. Fresh culture medium was added to quench the TrypLE and cells were resuspended and plated in new flasks and in fresh culture medium. At each passage, suspension cells were passaged by dilution into fresh culture medium in new flasks. The following cancer cell lines were purchased from the indicated vendors and cultured in the indicated media: 3T3 (CRL-1658) in DMEM, 293T (ATCC CRL-3216) in DMEM, U-2 OS (HTB-96) in DMEM, and K-562 (CCL-243) in Iscove's Modified Dulbecco's Medium (Sigma I3390). All cell cultures were maintained in an incubator at 37 °C with 5% $CO_2$ and humidity.

Engineered mammalian cell lines with MinDE circuits

Cell lines expressing MinDE circuits were generated for 3T3, 293T, U-2 OS, and K-562 cell lines. Lentiviral particles for each genetic construct were generated separately, then pooled, filtered to remove cellular debris, and concentrated to a final volume of 2 ml before addition. Constructs were expressed under the control of an EF1a promoter. Cells were sorted on an FACSAria cell sorter (BD Biosciences) based on MinD and MinE expression and subsequently expanded and frozen at low passage. Engineered cell lines were sub-cultured until cultures reached passage 35 at which point fresh cells were thawed. Cell lines with multi-part MinDE circuits were generated similarly or through lentiviral transduction of a single construct into an existing engineered cell line.

Engineered U-2 OS cell line with MinDE-Actuator circuit

U-2 OS cells were transduced with lentivirus to express the MinDE-Actuator circuit. Lentiviral particles for each genetic construct were generated separately, then pooled, supernatants were mixed, filtered to remove cellular debris, and concentrated to a final volume of 2 ml before addition. mCherry-MinD and MinE were expressed the control of an EF1a promoter, LifeAct-GFP was expressed under the control of a PGK promoter, and the Actuator construct was expressed under the control of a SFFV promoter. Cells were exposed to virus for 24 hours, then expanded for 24 hours in FluoroBrite™ DMEM (ThermoFisher A1896701) on glass bottom plates (Cellvis P06-1.5H-N) before immediately imaging.

Imaging MinDE circuits

Cells were plated on glass bottom plates (Cellvis P06-1.5H-N) to a confluency of 60-80% in FluoroBrite™ DMEM (ThermoFisher A1896701). After 1 day, cells were imaged on a Nikon Ti-Eclipse in a Tokai Hit stage-top incubator: 37°C and 5% $CO_2$ conditions. Confocal images were collected on a Nikon Ti-Eclipse equipped with a Yokogawa CSU-W1 confocal scanner unit in a Tokai Hit stage-top incubator: 37°C and 5% $CO_2$ conditions. Single and multi-channel fluorescence images were collected at 1 FPS. For long imaging experiments (typically over 2 hours), images were collected at 0.5 FPS.

Imaging experiments that required real-time stimulation with small molecules (rapamycin or isoprenaline) to activate MinDE circuits, were performed by direct addition of equal volume of media supplemented with 20μM of the respective small molecule into the well during image acquisition.

Image analysis

Basic image processing (cropping, background subtraction, and kymograph generation) was performed using Fiji ImageJ. Specific image-analysis workflows and frequency-domain signal processing was performed using custom python scripts as outlined below.

*Fast-Fourier Transform based frequency-domain image analysis*

Time-lapse fluorescence images were processed using fast-Fourier transform (FFT) to perform frequency-domain image analysis. Analysis was performed using custom written Python scripts. Time-lapse image data was processed by first zero-centering the individual pixel-level time-lapse data by subtracting polynomial decay. Pixel-level fluorescence intensity profiles were transformed from the time domain to the frequency domain using fast Fourier transform. Frequency bins 0 to N/2, where N is the number of time points sampled, were collected for each pixel to generate an image-level frequency domain spectrum. The absolute value of the image frequency spectra was taken to generate an image power spectrum. In this image power spectrum, cells expressing dynamic MinDE circuits will show oscillatory power in the frequency bin slice that corresponds to circuit frequency. Fiji ImageJ Temporal-Color Code was applied to the image power spectrum for visualization of cells expressing diverse MinDE circuit levels within a single image.

The image power spectrum served as the starting point for high-throughput frequency-domain image analysis. Individual frequency bin slices within the image power spectrum were treated as separate images. Each of these images were processed using standard OpenCV contour identification functions to identify and isolate the location of single cells that have oscillatory power within a particular frequency bin slice. The contour coordinates obtained were used as a reference to extract fluorescence intensity levels from the original time-lapse fluorescence image. The fluorescence intensities for all pixels within a contour were averaged across time then averaged with all pixels within the contour to calculate the average mCherry-MinD and MinE-GFP fluorescent intensity levels for a cell. The oscillatory power was calculated as the average of the power values within a contour in the respective frequency bin slice in the image power spectra. The frequency of the cell was assigned as the frequency bin slice in which it has maximum oscillatory power. Frequency bin numbers were converted to true frequency based on image acquisition parameters. This process was performed across the entire image power spectra to extract single-cell data within an image. Population wide MinDE circuit behaviors were analyzed by aggregating data across many images and thousands of cells.

*Hilbert Transform analysis of instantaneous MinDE circuit behavior*

Time-lapse fluorescence images were processed using Hilbert transform to report instantaneous MinDE circuit behavior. Analysis was performed using custom written Python scripts. Fast-Fourier transform was applied to the image to generate an image power spectrum. This spectrum was first inspected to determine which frequencies contain cells with oscillatory power. A finite impulse response (FIR) filter that isolated the frequencies of interest determined from the image power spectrum was applied to pixel-level fluorescence time-lapse data. This process removed contaminating frequencies from the time series image data and produced a filtered image with only MinDE signal content. Hilbert transform was applied to the filtered image to generate an analytic MinDE signal. The angle of the analytic signal produced a phase image series, and the absolute value of the analytic signal produced a power image series. These processed data reported on the instantaneous phase structure and instantaneous oscillatory power within an image. To generate a frequency image series, the pixel-level data in the phase image series were unwrapped to generate a continuous phase profile and the instantaneous derivative of this profile was the calculated instantaneous frequency. When necessary, FFT was used to identify frequency ranges associated with separate subcellular compartments and FIR filters were used to isolate these signals into separate images. The separate images were analyzed as described, and instantaneous oscillatory power image series were overlayed to visualize compartmentalization over time.

*Continuous Wavelet Transform analysis of instantaneous MinDE circuit behavior*

Complementary analysis of time-lapse fluorescence images using continuous wavelet transform (CWT) showed comparable results to Hilbert transform. Analysis was performed using custom written Python scripts. The ssqueezepy Python package (pypi.org/project/ssqueezepy/) was utilized to generate synchrosqueezed scalograms. CWT was applied directly to pixel-level fluorescence time series to generate pixel-level scalograms. The absolute value of the pixel-level scalograms were taken and the frequency for the pixel was assigned as the frequency corresponding to the scale with maximum magnitude. The amplitude for the pixel was assigned as the magnitude at that scale frequency and the phase was assigned as the angle of the original scale coefficient at the scale frequency. This was done for all time points of the pixel-level scalogram for all pixels within the image to generate instantaneous phase, frequency, and amplitude image series using CWT.

<u>Spatial analysis using instantaneous phase image series</u>

Time-lapse fluorescence images were transformed into phase image series using Hilbert transform. Standard OpenCV optical flow functions were applied to sequential images in the phase image series to generate vector fields describing pixel-level spatial movement of patterns between time intervals. The Runge-Kutta Fourth Order method for solving the initial value problem for partial differential equations was used to draw a vector-field guided path that described the spatial movement of a pattern from a single pixel. The phase values along the vector-guided path were collected, and the derivative of the continuous phase along the path per $2\pi$ represents the spatial wavelength of the cell.

Custom written Python scripts were used to analyze locations of spiral centers within cells in images. First, enclosed square paths were generated for every pixel within an image. Then, phase values in the Hilbert transform phase image series were collected according to the path and the sequential sum of these values represented the vorticity. The values in the vorticity image near $2\pi$ and $-2\pi$ represented the spiral centers. This was applied to every frame within the phase image series to generate a spiral center image series. The sign of the vortex determined it's clockwise or counterclockwise direction of rotation.

<u>High-throughput PKA dynamics reporter analysis</u>

Time-lapse fluorescence images for the MinD and FHA1 channels were subjected to a high-pass FIR filter to zero center pixel-level fluorescence intensity time profile. The Hilbert transform analysis pipeline was applied as described to generate instantaneous power image series for each respective channel. Pixel-level ratios of FHA1 power to MinD power were performed to normalize fluctuations in power. Power normalization generated an instantaneous PKA dynamics image series that reported changes in PKA activity over time. A low-pass FIR filter was applied to this image to filter out fluctuations.

Frequency-based cell segmentation was applied to the filtered image series to locate and isolate cells expressing MinDE circuits. The locations of the cells were used to extract PKA activity dynamics from the power normalized image series. Pixel-level profiles for all pixels corresponding to a cell were averaged to assign the cell's PKA signaling trajectory. PKA signaling trajectories for all cells were collected and hierarchical clustering was applied to the time profiles to group cells with similar PKA activity dynamics.

Cross-correlation analysis of MinD-ActA to LifeAct in actin patterning circuits

Time-lapse fluorescence images for the MinD and LifeAct channels were subjected to a high-pass FIR filter to zero center pixel-level fluorescence intensity time profile. For each pixel, the cross-correlation between the LifeAct and MinD signals (post rapamycin induced coupling) was computed to create a profile representing the correlation of the two signals as a function of time-lag (see Fig. 4I or S11). The best time lag at each pixel in the image was then extracted by taking the argmax of the resulting cross-correlation profile. The resulting image-scale representation was used to display and inspect how the time-lag between the two signals was distributed throughout the cell.

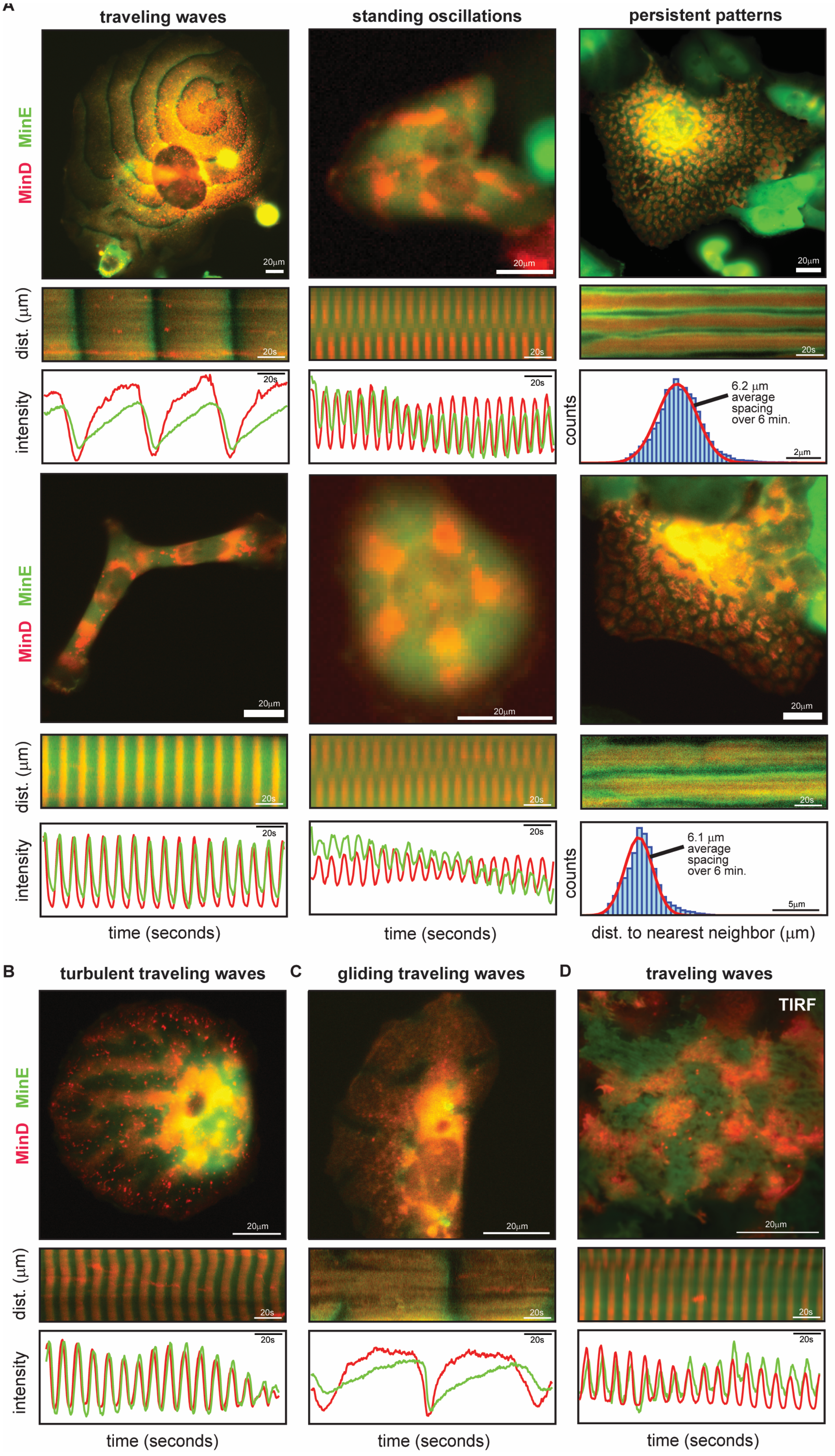
A
traveling waves
standing oscillations
persistent patterns
MinD MinE
20μm
dist. (μm)
20s
intensity
counts
6.2 μm average spacing over 6 min.
2μm
6.1 μm average spacing over 6 min.
5μm
time (seconds)
dist. to nearest neighbor (μm)
B
turbulent traveling waves
C
gliding traveling waves
D
traveling waves
TIRF

**Figure S1. MinDE reaction-diffusion system generates diverse spatiotemporal protein patterns in mammalian cells.**

(A) MinDE reaction-diffusion system generates subcellular traveling waves, standing oscillations, and persistent "leopard print" patterns of protein localization in metazoan cells. Composite epifluorescence images of mCherry-MinD and MinE-EGFP show instantaneous subcellular MinDE patterns. Kymographs show differences in spatiotemporal behavior between different reaction-diffusion patterns. Traveling wave kymographs show uniform temporal spacing between periodic wavefronts. Kymographs of slow traveling waves enable visualization of MinE-EGFP lag at the edge of the wavefront. Standing oscillation kymographs show alternating pairwise periodic nodal structures across time. Persistent "leopard print" pattern kymographs show spots are stable across time. Pixel-level data for traveling waves and standing oscillations show strong oscillatory nature of MinDE spatiotemporal patterns. All pixel-level data are zero-centered and normalized to show colocalization between mCherry-MinD and MinE-EGFP signals. Histogram of distances between nearest neighbor spots shows persistent patterns are systematically spatially organized. See also Movie S3-5.

(B) Traveling wave patterns within a single cell may have more than one spiral center. This results in turbulent patterns where many waves crash and combine with each other at various locations within the cell. Composite image level representation of these patterns shows chaotic nature of MinDE reaction-diffusion behavior in these cells. Kymographs and pixel-level analysis of these chaotic patterns show that the MinDE reaction-diffusion in these cells still obey oscillatory dynamics seen in simple traveling wave patterns. See also Movie S6.

(C) Traveling wave patterns in cells expressing high amounts of mCherry-MinD relative MinE-EGFP are characterized by "negative" wavefronts. "Positive" wavefronts are described as the movement of mCherry-MinD and MinE-EGFP in the wave, while "negative" wavefronts are described as the depletion of mCherry-MinD and MinE-EGFP in an area by the wave. Kymographs and pixel-level representations show that these patterns obey oscillatory dynamics seen in other traveling waves, however, have extremely slow oscillations. See also Movie S6.

(D) TIRF images show traveling wave patterns in metazoan cells. See also Movie S6.

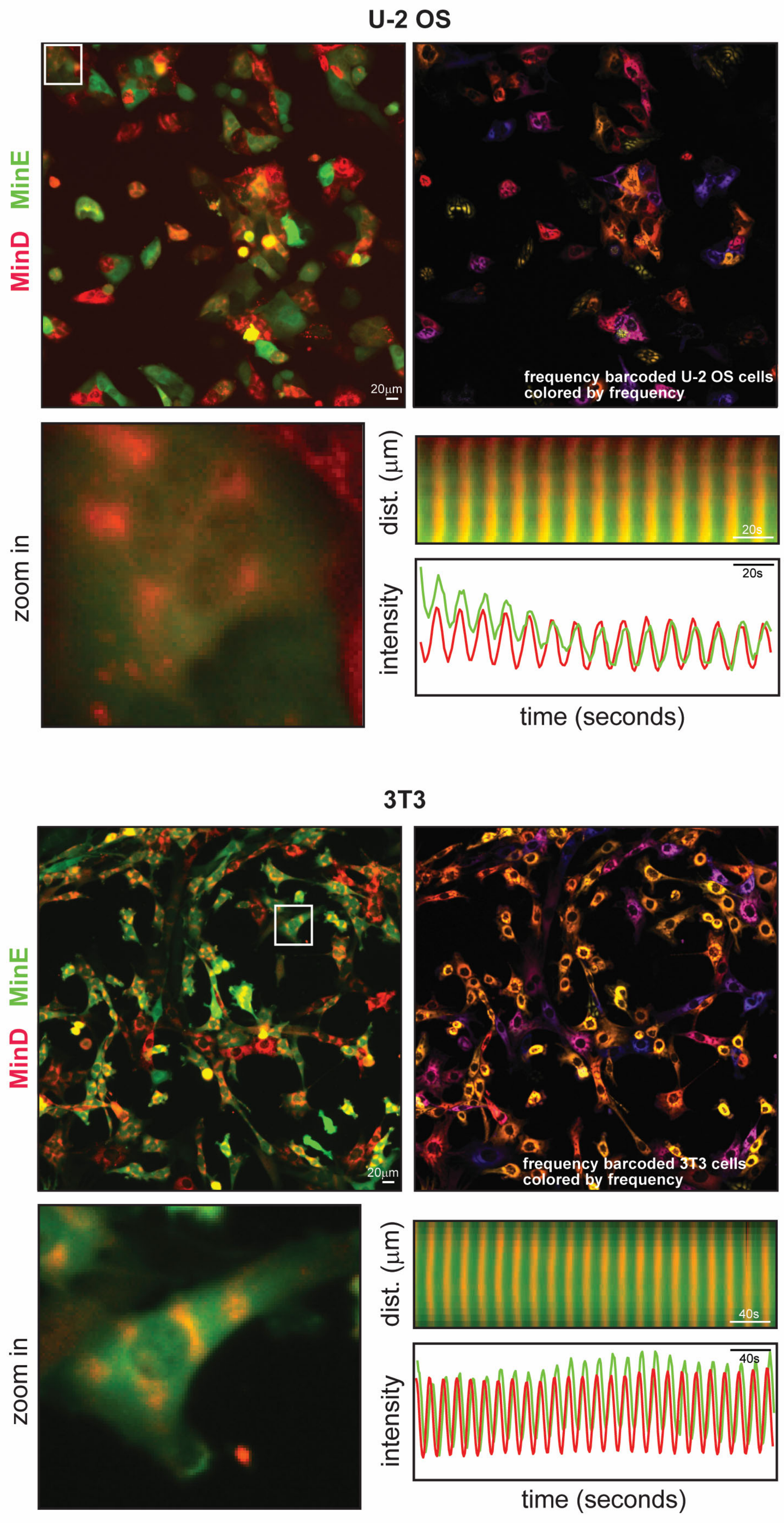
U-2 OS
MinD MinE
20μm
frequency barcoded U-2 OS cells
colored by frequency
zoom in
dist. (μm)
20s
intensity
20s
time (seconds)
3T3
MinD MinE
20μm
frequency barcoded 3T3 cells
colored by frequency
zoom in
dist. (μm)
40s
intensity
40s
time (seconds)

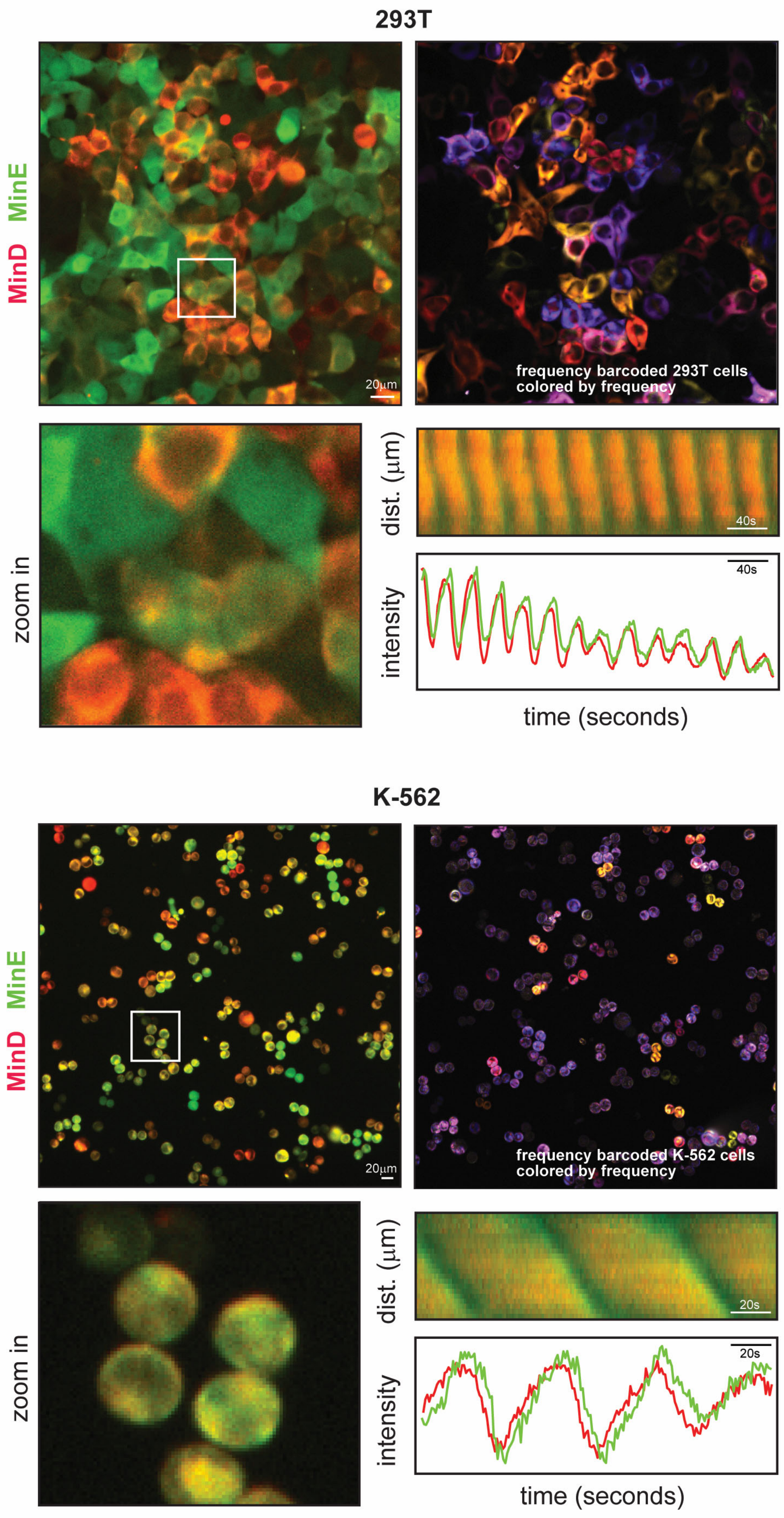

293T
MinD MinE
20μm
frequency barcoded 293T cells colored by frequency
zoom in
dist. (μm)
40s
intensity
40s
time (seconds)
K-562
MinD MinE
20μm
frequency barcoded K-562 cells colored by frequency
zoom in
dist. (μm)
20s
intensity
20s
time (seconds)

**Figure S2. MinDE circuits have activity in a broad range of cell types.**

MinDE circuits are active when installed in adherent (U-2 OS, 3T3, and 293T) and suspension (K-562) cell lines. Low-magnification composite mCherry-MinD and MinE-EGFP epifluorescence images of different cell types expressing MinDE circuits show stable cell lines that display active MinDE circuits can be generated for a variety of cell types. Application of frequency-domain image analysis enables frequency-domain cell segmentation and FM-barcoding of individual cells regardless of cell type or cell shape (Fig. 2). Manual analysis (kymograph and pixel-level data) of single cells within the field of view for each cell type show that MinDE reaction-diffusion traveling waves show consistent behavior across cell types. See also Movie S12.

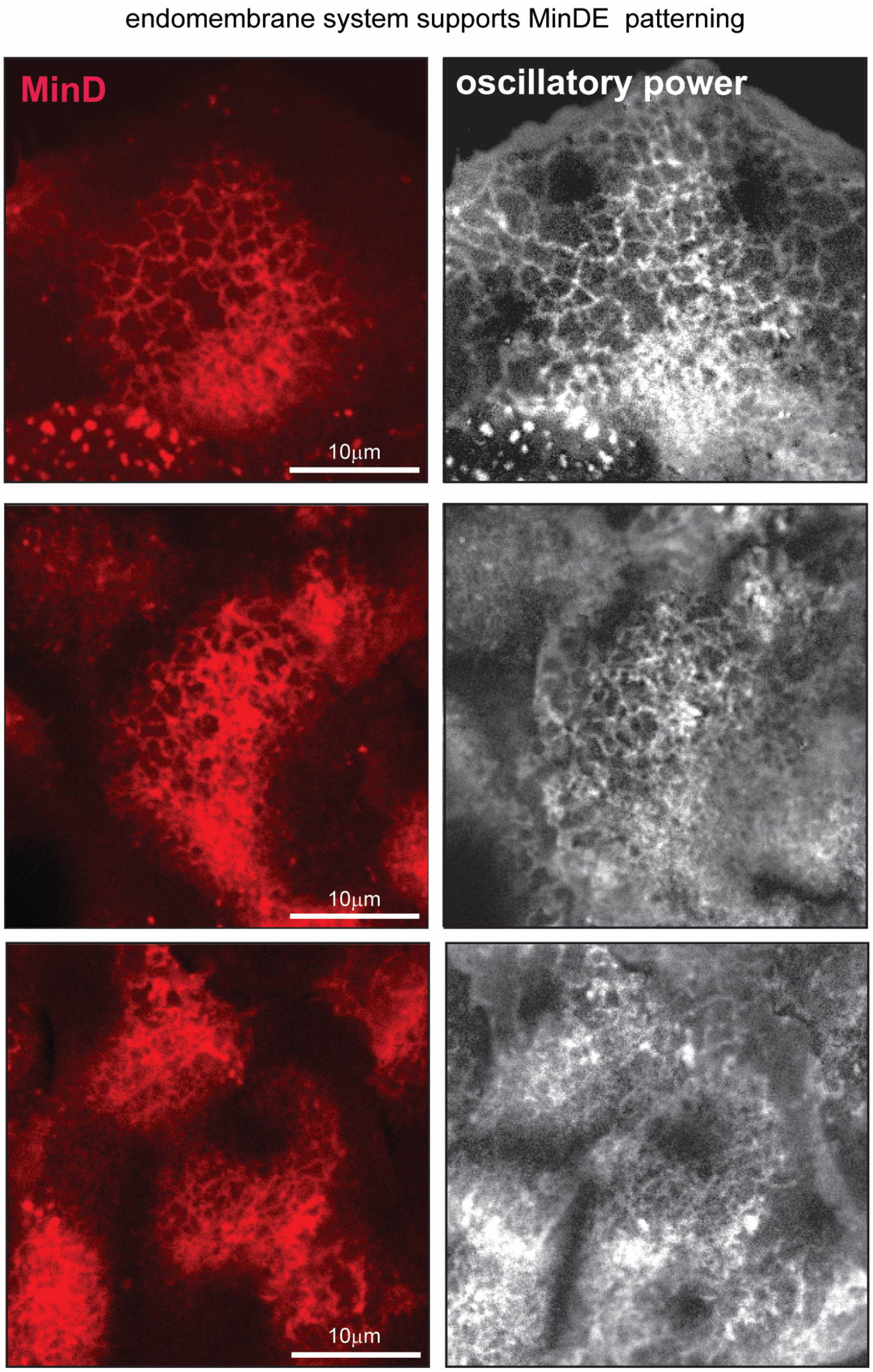


**Figure S3. Confocal imaging suggests MinDE patterning is supported by the endomembrane system.**

High-magnification confocal fluorescence imaging of the mCherry-MinD signal within mammalian cells shows that MinDE patterns are colocalized to the endomembrane system. FIR (finite impulse response) filters were applied to these confocal image time series data to isolate oscillatory content in the data. Hilbert transform was applied to the FIR filtered images to extract the oscillatory power (Hilbert transform envelope) of different regions within the cell (Fig. S4). Oscillatory power image representations of the confocal fluorescence images show that oscillatory power is concentrated in regions colocalized to the endomembrane system. See also Movie S7.

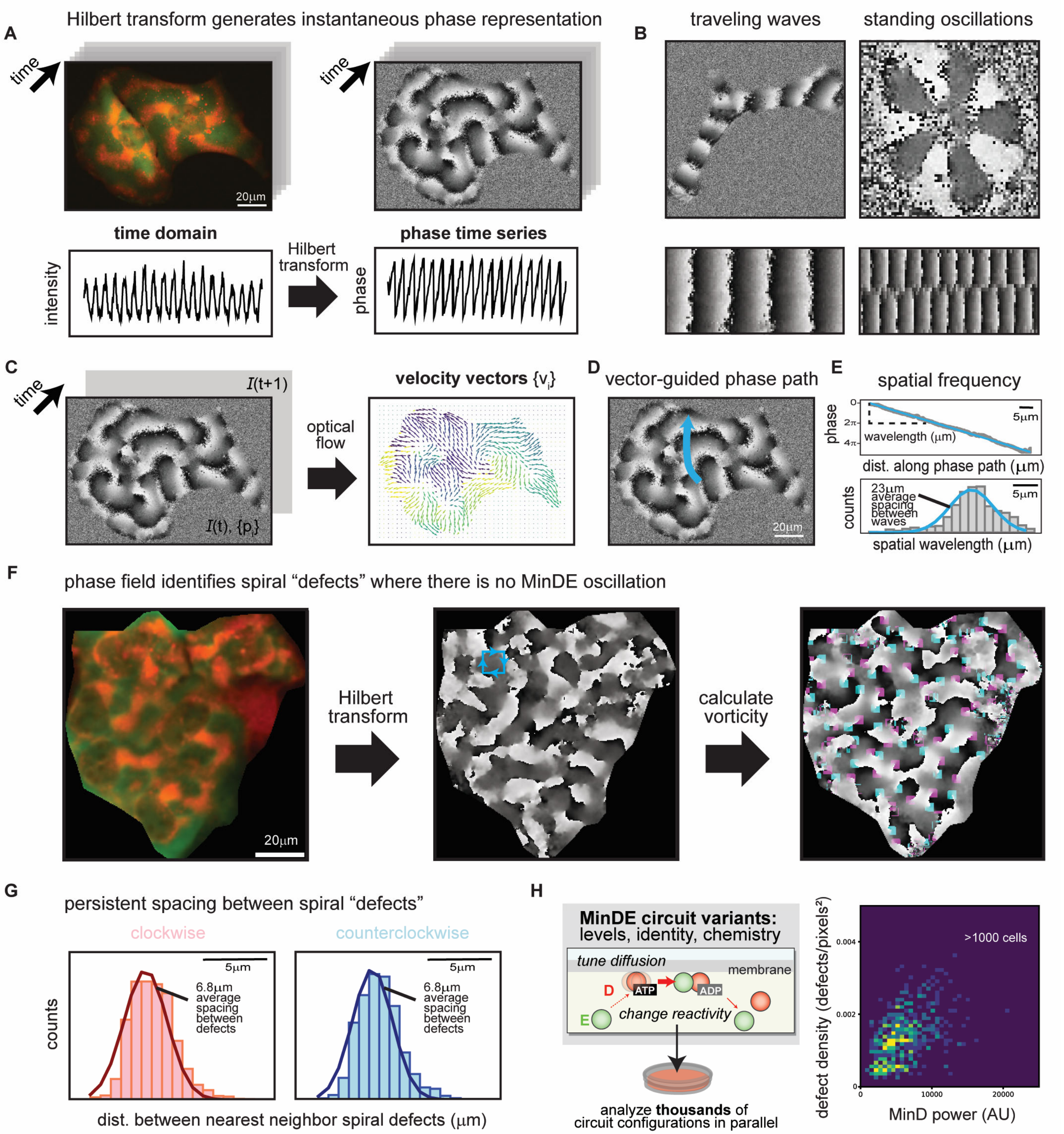


**Figure S4. Hilbert transform derived phase-field representations enable spatial characterization of MinDE dynamic patterning.**

(A) Hilbert transform generates phase image time series (Fig. S4B).

(B) Phase image representations of dynamic oscillatory MinDE patterns enable characterization of traveling waves and standing oscillations. Image-level inspection of phase images shows traveling wave patterns display a smooth gradient of phase angles from 0 to 2π between wavefronts while standing oscillations display pairwise bands of two different phase angles. The kymograph of the phase image time series for the travel waves again shows a smooth gradient of phase angles from 0 to 2π between wavefronts. The kymograph for standing oscillation shows a pairwise alternating nodal structure across time. See also Movie S8.

(C) The phase image time series can be used to generate velocity maps that report the spatial movement of patterns across time. Standard OpenCV optical flow algorithms can be applied to two sequential frames in the phase image time series to generate a vector field that describes the spatial direction in which the pattern will move for every pixel within an image. FFT-based frequency domain analysis (Fig. 2) can be used to identify, isolate, and FM-barcode cells within the image to identify the regions containing cells of interest to extract vector field information for only those cells and to assign that information to those cells. This enables vector map representation that describes spatial MinDE circuit behavior within that cell across time.

(D) Following the path taken by the pattern using the vector map representation for each instant in time allows for calculation of spatial frequency within the cell. The Runge-Kutta method was used to draw vector map guided paths within a cell when starting from a signal pixel.

(E) Spatial characterization of MinDE circuit behavior was performed by combining the phase field image with the associated vector map. After determining the path from a starting pixel (Fig. S5D), the phase values along that path were extracted from the phase image. The slope of the unwrapped phase angle values to distance across vector-guided path per $2\pi$ is defined as the spatial wavelength of the cell's pattern. This was repeated using several hundred unique starting pixels to generate a data set containing many spatial wavelengths for a cell of interest. This data set supports that there is consistent spacing between wavefronts and that dynamic oscillatory MinDE signals are not only temporally organized, but also systematically spatially organized.

(F) Phase field representations of MinDE image signals can be used to locate spiral centers or "defects" within the cell. These are regions within the cell where there is no MinDE oscillation as a result of being locations where waves originate or crash and dissipate. To calculate vorticity, for each pixel, a square vortex path that starts and ends at that starting pixel was generated. The phase field values along the direction of the path were collected and the sequential sum of the path reported the vorticity for the pixel. Regions where the vorticity is approximately $2\pi$ or $-2\pi$ are spiral centers/defects. The sign of the vorticity value indicates the directionality of the rotation. The spiral centers are displayed on the phase field and are color-coded to show clockwise (pink), and counterclockwise (blue) spiral centers located with the cell. See also Movie S9.

(G) Analysis of nearest neighbor distances between spiral centers shows that there is consistent spacing between spiral centers that share the same direction of rotation and within the same cell, both positive-positive and negative-negative spiral centers are spaced the same distance away.

(H) High-throughput analysis of single-cell data for thousands of cells (U-2 OS) shows biochemical trends govern the defects density found within a cell. A 2D histogram relating defect density (total area of defects per total area of cell) to MinDE power indicates a positive correlation between MinDE power levels and defect density suggesting that cells expressing MinDE circuit configurations with high power will display turbulent patterning phenomena.

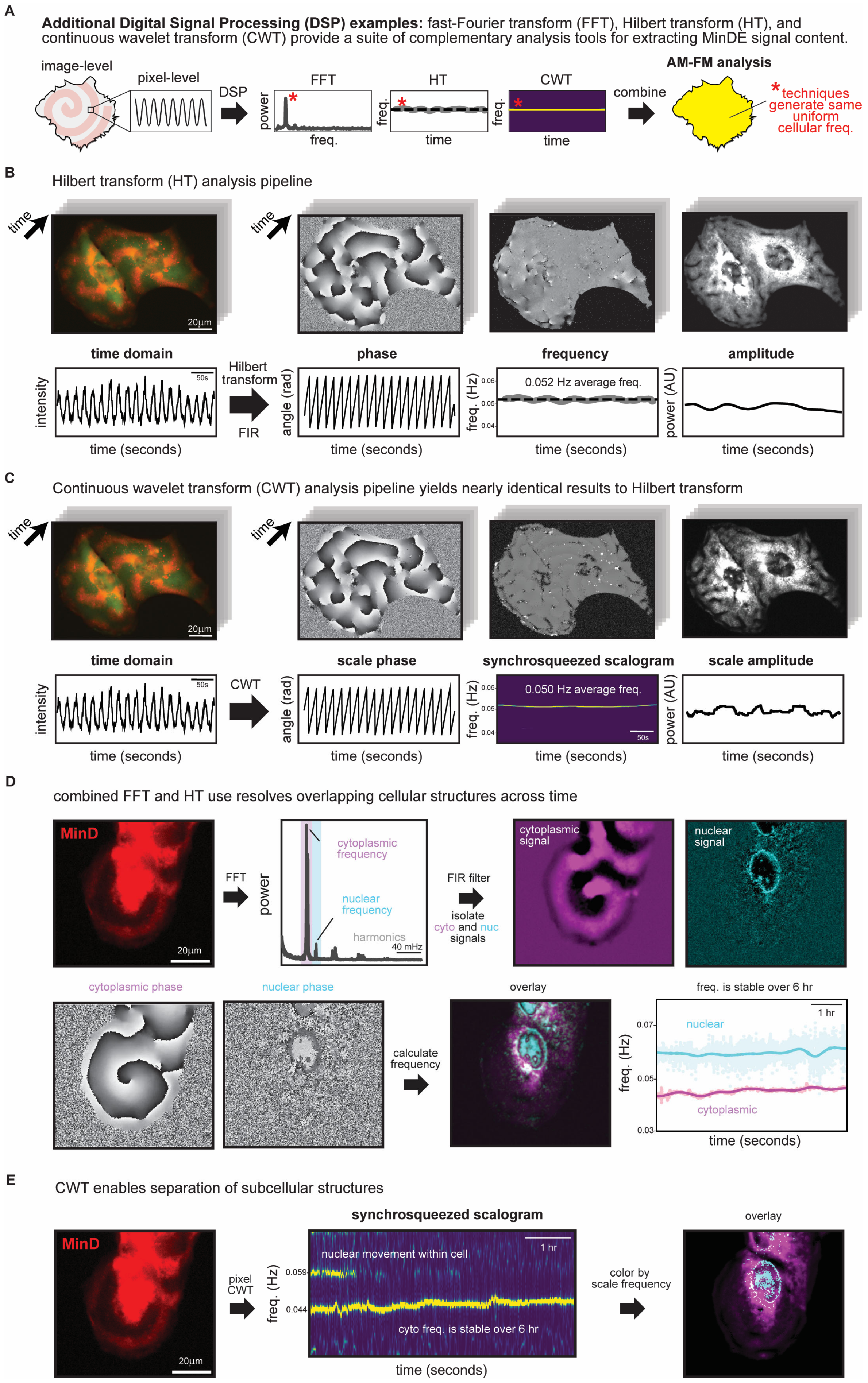

A
Additional Digital Signal Processing (DSP) examples: fast-Fourier transform (FFT), Hilbert transform (HT), and continuous wavelet transform (CWT) provide a suite of complementary analysis tools for extracting MinDE signal content.
image-level
pixel-level
DSP
FFT
power
freq.
HT
freq.
time
CWT
freq.
time
combine
AM-FM analysis
* techniques generate same uniform cellular freq.
B
Hilbert transform (HT) analysis pipeline
time
20µm
time
time domain
phase
frequency
amplitude
intensity
50s
time (seconds)
Hilbert transform
FIR
angle (rad)
time (seconds)
freq. (Hz)
0.052 Hz average freq.
0.06
0.05
0.04
time (seconds)
power (AU)
time (seconds)
C
Continuous wavelet transform (CWT) analysis pipeline yields nearly identical results to Hilbert transform
time
20µm
time
time domain
scale phase
synchrosqueezed scalogram
scale amplitude
intensity
50s
time (seconds)
CWT
angle (rad)
time (seconds)
freq. (Hz)
0.050 Hz average freq.
0.06
0.05
0.04
50s
time (seconds)
power (AU)
time (seconds)
D
combined FFT and HT use resolves overlapping cellular structures across time
MinD
20µm
FFT
power
cytoplasmic frequency
nuclear frequency
harmonics
40 mHz
FIR filter
isolate cyto and nuc signals
cytoplasmic signal
nuclear signal
cytoplasmic phase
nuclear phase
calculate frequency
overlay
freq. is stable over 6 hr
1 hr
nuclear
cytoplasmic
freq. (Hz)
0.07
0.05
0.03
time (seconds)
E
CWT enables separation of subcellular structures
MinD
20µm
pixel CWT
synchrosqueezed scalogram
freq. (Hz)
0.059
0.044
nuclear movement within cell
1 hr
cyto freq. is stable over 6 hr
time (seconds)
color by scale frequency
overlay

**Figure S5. Digital Signal Processing techniques enable characterization of instantaneous MinDE circuit behavior.**

(A) Fluorescence microscopy of MinDE circuit behaviors in mammalian cells generates images encoding dynamic oscillatory signals that describe the reaction-diffusion patterning phenomena. These oscillatory signals encoded within fluorescence image time series data can be analyzed using Digital Signal Processing (DSP) techniques such as fast-Fourier transform (FFT), Hilbert transform (HT), and continuous wavelet transform (CWT) to extract MinDE signal content. Pixel-level analysis of fluorescence image data using DSP allows for extraction of frequency information content across an entire image. The FFT, HT, and CWT provide a suite of complementary analysis tools to locate, isolate, and analyze frequency information content for downstream applications such as frequency-domain image processing and oscillatory power-based host-cell activity reporting.

(B) Hilbert transform enables characterization of instantaneous MinDE signal content across time. The FFT analysis pipeline is applied to image-level data (Fig. 2B) to locate the frequencies in the frequency-domain where a cell's MinDE circuit has oscillatory power. A narrow-band FIR filter is applied to the image around the frequencies located in the FFT to isolate the cell's frequency signal content within the fluorescence image time series. The HT is applied to the filtered image to generate an analytic MinDE time signal. The angle of the analytic signal generates the instantaneous phase representation, and the magnitude of the analytic signal generates the instantaneous amplitude representation. The phase imparts information on the momentary structure of the MinDE pattern in an instant in time while the amplitude imparts information on the oscillatory power. The instantaneous frequency is generated by taking the derivative of the unwrapped phase time signals. See also Movie S14.

(C) Continuous wavelet transform yields nearly identical results to the Hilbert transform. CWT is applied directly to fluorescence image times series data. This generates pixel-level scalograms reporting on frequency power content across times for different scales that each represent a unique frequency. A CWT package that utilizes synchro-squeezing in the scalogram representation of the oscillatory signal data was used. The scale with the largest magnitude represents the strongest oscillatory frequency detected within a pixel-level MinDE signal and is assigned as the pixel's frequency. The magnitude of scale coefficient itself represents the oscillatory power and the angle of the scale coefficient is the phase. HT and CWT analysis of the same cell report nearly identical phase, frequency, and oscillatory power information. See also Movie S16.

(D) Strategic use of FFT, FIR filters, and HT enables instantaneous characterization of subcellular structures across time. FFT is used to locate the signals of interest, FIR filters are used to isolate these signals from the fluorescence image time series data, and these signals are analyzed separately using Hilbert transform to extract instantaneous frequency content. The analysis pipeline shows how cytoplasmic and nuclear signals can be simultaneously tracked across long periods of time. This enables visualization of subcellular organelle location using single-channel fluorescence microscopy data. See also Movie S15.

(E) Similar to analysis pipeline discussed in Fig. S4D, CWT enables separation and visualization of subcellular structures. However, the CWT does not require pre-processing to achieve this. CWT separates cytoplasmic and nuclear signals into different scale frequencies across the entire duration of the time series data. By visualizing the different scale frequencies that contain power for signals of interest, it is possible to directly isolate cytoplasmic and nuclear signal content. See also Movie S17.

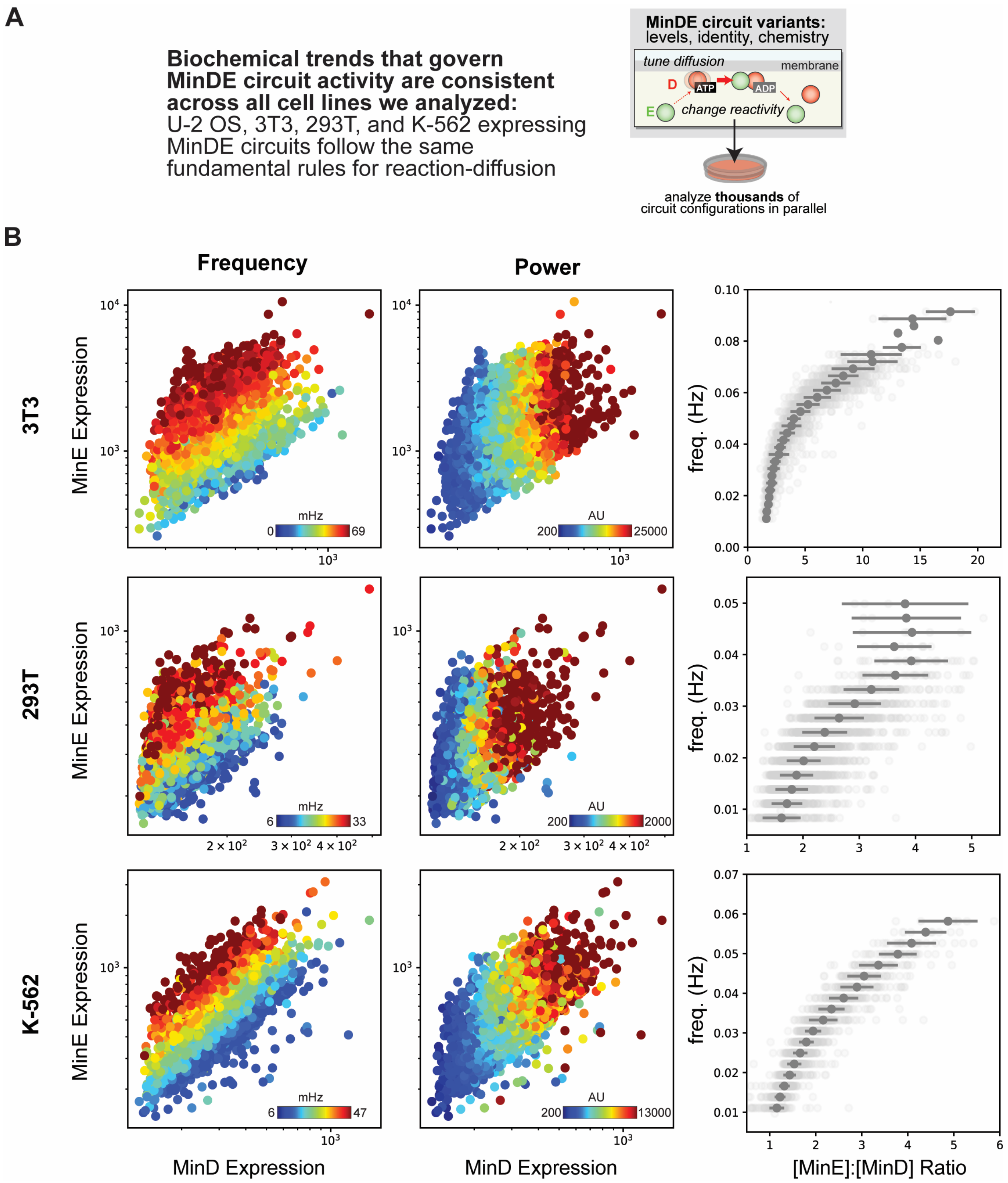


**Figure S6. Biochemical trends governing MinDE circuit behaviors are consistent across a broad range of cell types.**

(A) High-throughput analysis of single-cell data for thousands of cells per cell type (3T3, 293T, and K-562) was used to show biochemical trends in MinDE circuit behavior. For each cell type, many low-magnification epifluorescence images of mCherry-MinD and MinE-GFP (Fig. S2) were collected. Each image collected data on hundreds of cells displaying active MinDE circuit behavior. The frequency-domain analysis workflow was used to spectrally separate, segment, and FM-barcode each cell in an epifluorescence image time series. Standard OpenCV image processing strategies were used to isolate FM-barcoded cells to extract oscillatory frequency, power, and fluorescent intensity levels for each cell (fluorescence intensity levels are used as a proxy for protein expression levels). Single cell data extracted across many low-magnification fields of views were aggregated to generate data sets containing information on thousands of cells for each cell type.

(B) The data across the cell types show that the frequency of dynamic MinDE patterns is governed by MinE expression levels: as MinE expression levels increase, frequency increases. Oscillatory power of dynamic MinDE patterns is governed by MinD expression levels: as MinD expression levels increase, oscillatory power increases. Patterning frequency is governed by the ratio of MinE to MinD in the cell: as the ratio of MinE to MinD increases, frequency increases. The consistency of these relationships across cell types shows there is a standard set of biochemical rules that govern the behavior of the MinDE reaction-diffusion system in mammalian cells.

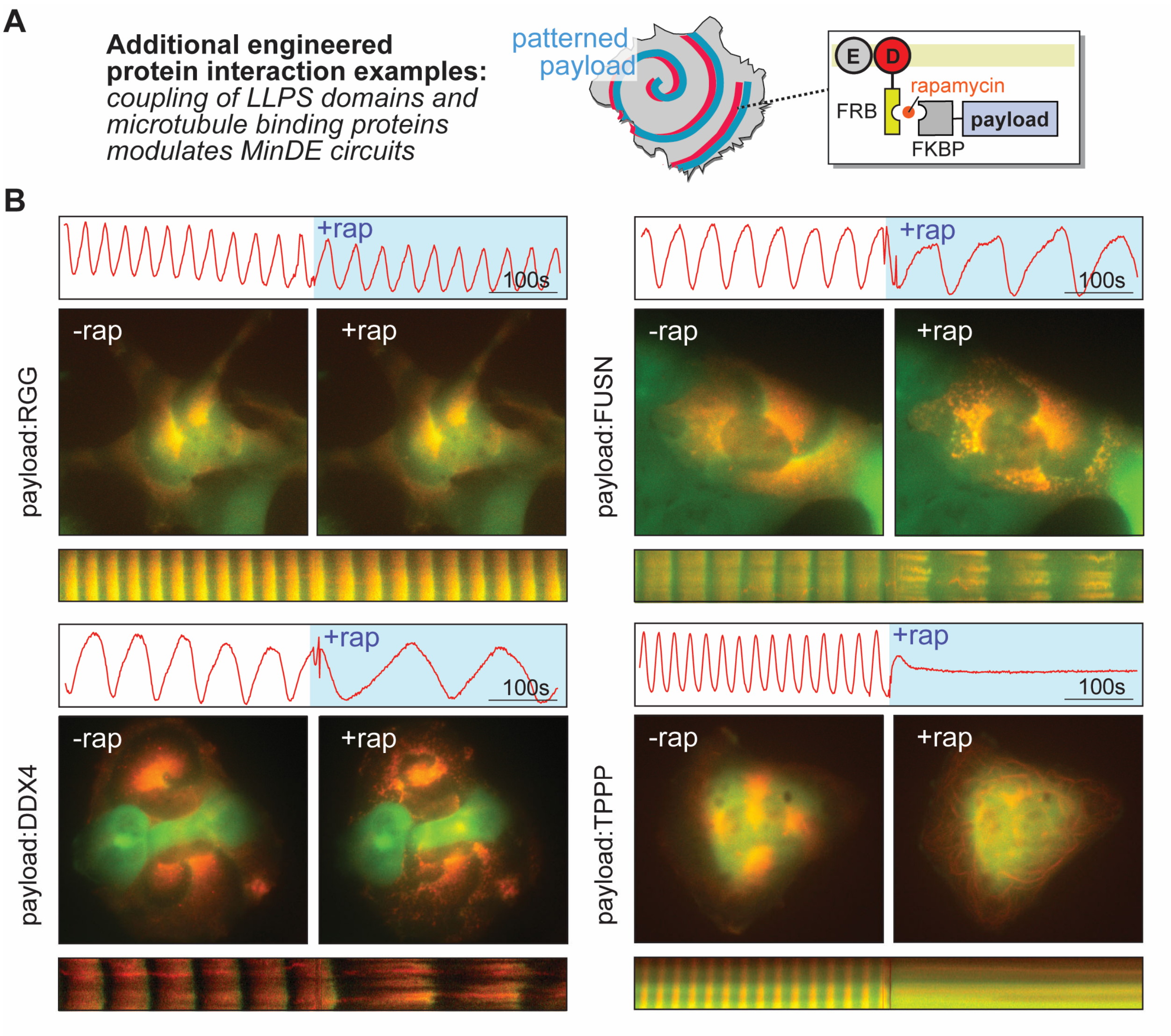


**Figure S7. Recruitment of protein payloads can dynamically modulate MinDE reaction-diffusion behavior.**

(A) The baseline MinDE circuit behavior is genetically encoded and governed by biochemical conditions within the cell (Fig. S6). Engineered protein interactions enable real-time modulation of MinDE reaction-diffusion behavior by dynamically altering system conditions. The chemical inducible dimerization (CID) pair FRB-FKBP is used to controllably load protein payloads with high specificity and affinity onto pre-existing MinDE patterning phenomena. See also Movie S18.

(B) RGG (Arginine-Glycine-Glycine), FUS (Fused in Sarcoma), and DDX4 (DEAD-box helicase 4) are LLPS (liquid-liquid phase separation) modules with varied multivalent macromolecular interaction strength (RGG < FUSN < DDX4). Each LLPS module interacts with other copies of itself to form phase condensates, membrane-less organelles with high protein density and reduced diffusivity to the surrounding environment. Loading the LLPS module onto the MinDE circuit causes real-time frequency modulation of reaction-diffusion behavior where decrease in frequency is proportional to the strength of the LLPS modules (seen in the kymograph and pixel-level intensity time profile). Composite epifluorescence images of patterning phenomena show upon coupling, MinDE proteins cluster into protein condensates and these condensates are still spatiotemporally organized into traveling waves within cell. TPPP (Tubulin polymerization-promoting protein) binds to microtubules. Loading TPPP onto the MinDE circuit sequesters MinD to the cytoskeletal microtubules and abolishes reaction-diffusion behaviors (seen in the kymograph and pixel-level intensity time profile). See also Movie S19.

**DSP Workflow for extracting FM-barcoded single-cell PKA signaling dynamics trajectories.**

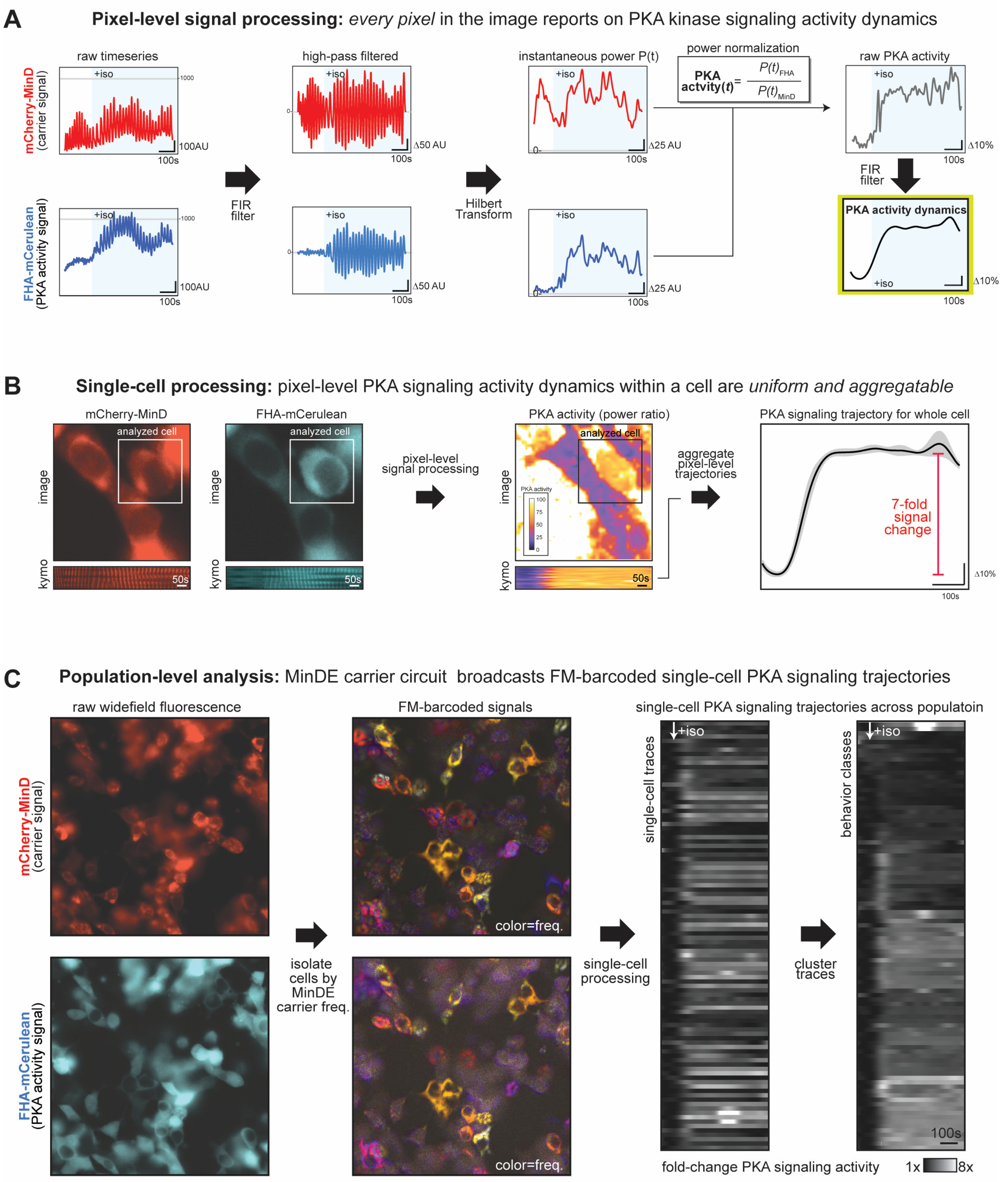

**Figure S8. Digital signal processing workflow for extracting FM-barcoded single-cell PKA signaling dynamics trajectories**

PKA signaling trajectories were computed at a single-pixel level and aggregated at the cellular level based on FM-barcode using the frequency-domain cell segmentation strategy outline in the supplemental methods. The workflow for each step is illustrated below. See also Movie S20-21.

(A) At the single-pixel level, the raw fluorescence time series for the MinD (carrier line) and FHA-mCerulean (PKA data line) show that the FHA signal begins co-oscillating with the MinD signal upon addition of isoprenaline. The DC components of this signal were removed using an FIR filter, resulting in zero-centered signals with identical frequencies but different time-dependent amplitudes. The instantaneous power throughout the time course was calculated using the Hilbert Transform. Because the FHA signal is dependent on association with MinD for its oscillation, the instantaneous power of the MinD signal can be used to normalize the instantaneous power of the FHA signal against spurious fluctuations. The resulting signal, *PKA-activity(t)*, shows a large increase to a new steady-state value about isoprenaline addition. High-frequency fluctuations in this profile can then be removed by applying a low-pass FIR filter.

(B) To obtain single-cell PKA signaling trajectories, the pixel-level data in (A) was binned across all pixels belonging to an individual cell, where cell boundaries were located using FM-barcodes and the frequency-domain cell-segmentation workflow in (C). Because the pixel-level PKA signaling data is normalized to MinD signal, the resulting trajectories are similar everywhere within the cell, even when intensities vary owing to differences to the thinness of the cell. This allows the single-pixel trajectories to be averaged to produce a single-cell trajectory along with its variance. For the example cell shown, the addition of isoprenaline produces a smooth PKA-activity trajectory that levels off to a new steady state after approximately 100 seconds. For the indicated cell, the measurable change in PKA-activity was 7-fold higher after stimulation.

(C) To extract multiple single-cell trajectories from wide-field imaging of a population of cells, frequency-domain cell-segmentation was performed using the MinD signal as described in the supplemental methods. Inspection of FM barcodes for the MinD (carrier signal) and FHA-mCerulean (PKA activity signal) shows that the FHA-mCerulean signals oscillate at the same frequency as the MinD carrier. The PKA signaling activity trajectory for 98 different cells was extracted as described in (B) and clustered to reveal trends in the associated responses. Using this method, most cells showed an increase to a new-steady state level of PKA signaling over the span of approximately 100 seconds.

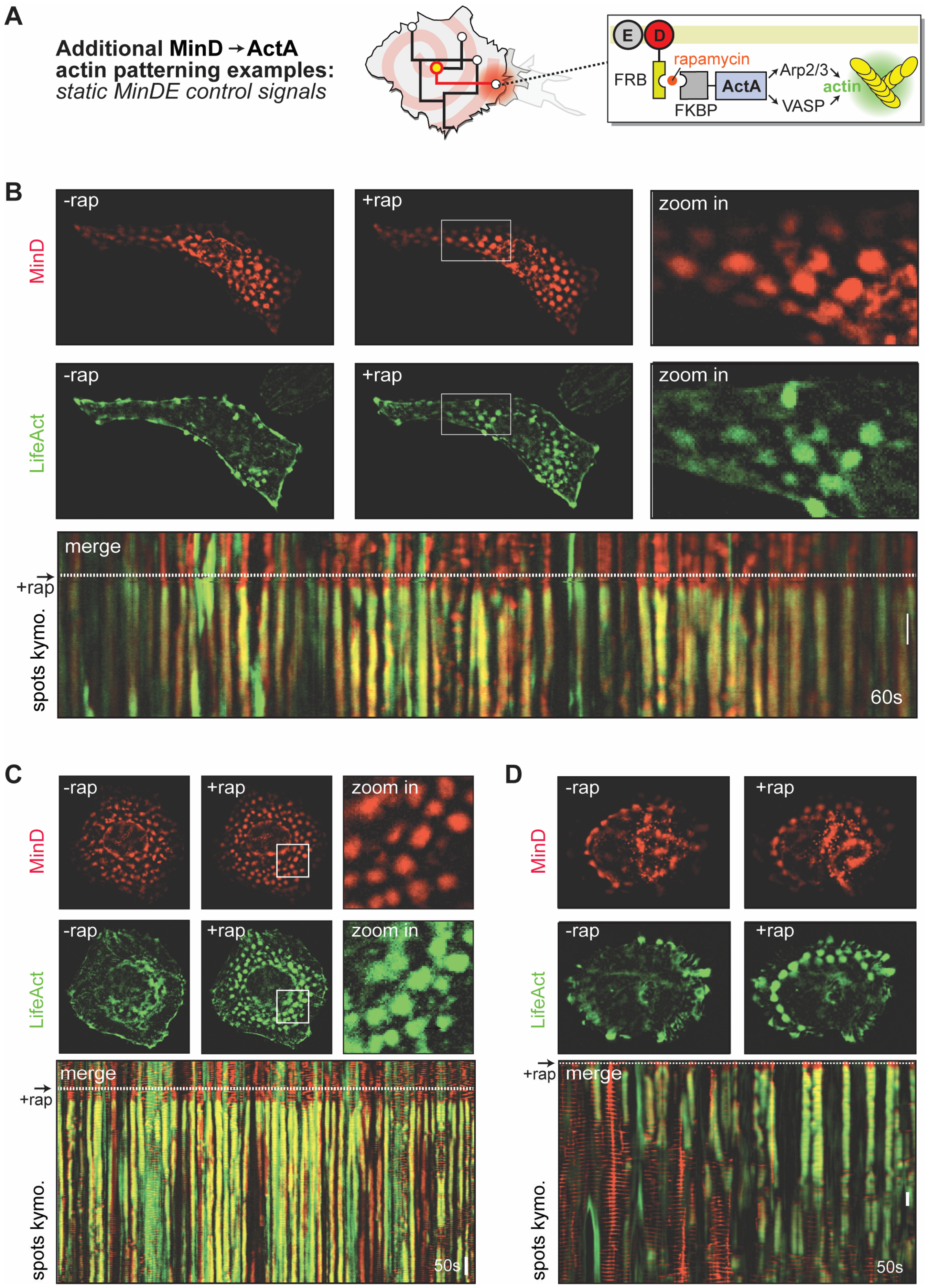

A
Additional MinD→ActA
actin patterning examples:
static MinDE control signals
E
D
FRB
rapamycin
FKBP
ActA
Arp2/3
VASP
actin
B
MinD
-rap
+rap
zoom in
LifeAct
-rap
+rap
zoom in
merge
+rap
spots kymo.
60s
C
MinD
-rap
+rap
zoom in
LifeAct
-rap
+rap
zoom in
merge
+rap
spots kymo.
50s
D
MinD
-rap
+rap
LifeAct
-rap
+rap
+rap
merge
spots kymo.
50s

**Figure S9. Additional examples of MinD-ActA driven actin patterning using static MinDE control signals.**

(A) Additional examples of MinDE to ActA patterning circuits in which the MinDE control signal is static, forming spotted regions of MinD within the cell that are stationary over an extended period. These control signals are inducibly connected to the ActA effector protein upon addition of rapamycin, which will stimulate actin polymerization through recruitment of Arp2/3 and VASP.

(B) and (C) Examples of spotted actin polymerization templated by the MinDE-ActA circuit. Representative images of MinD or the LifeAct reporter, pre and post rapamycin addition are shown. A kymograph drawn through the spotted pattern of the MinD and LifeAct signals is also shown. See also Movie S22-23.

(D) As in (B) and (C), but for MinDE control signal that is less stable and less persistent over time. This results in more stable actin structures as judged by the LifeAct signal in the kymograph. See also Movie S22-23.

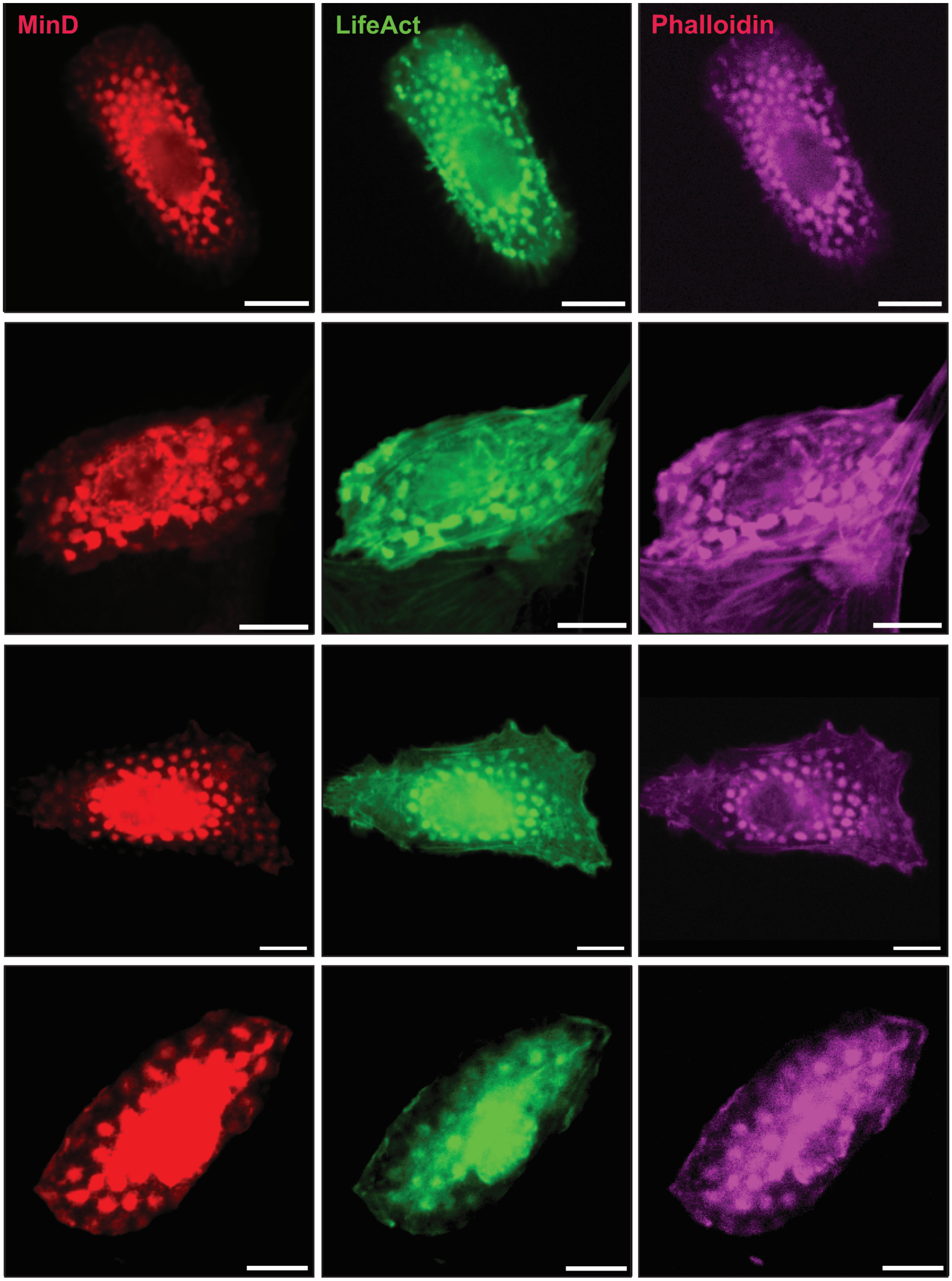


**Figure S10. Phalloidin staining confirms MinDE-ActA circuit directed assembly of synthetic filamentous actin structure.**

Cells harboring MinDE-ActA circuits were stimulated with rapamycin, fixed and permeabilized, and stained for filamentous actin using phalloidin. Phalloidin staining colocalized with fixed MinD “leopard print” patterns and agrees with LifeAct reporter signal for actin polymerization. (scale bar: 20μM)

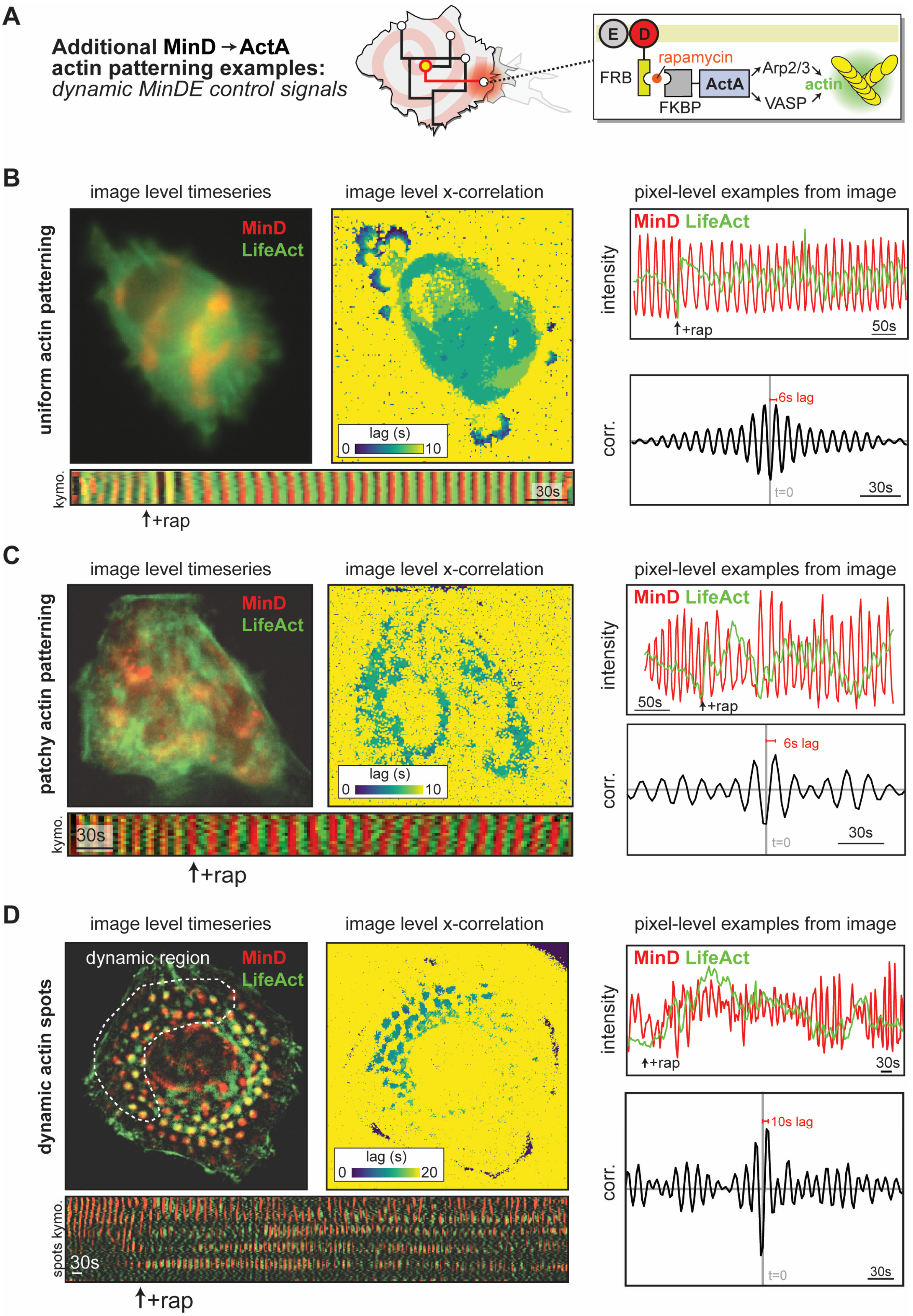
A
Additional MinD → ActA
actin patterning examples:
dynamic MinDE control signals
E
D
FRB
rapamycin
FKBP
ActA
Arp2/3
VASP
actin
B
image level timeseries
image level x-correlation
pixel-level examples from image
uniform actin patterning
MinD
LifeAct
lag (s)
0
10
kymo.
30s
+rap
intensity
50s
6s lag
corr.
t=0
30s
C
image level timeseries
image level x-correlation
pixel-level examples from image
patchy actin patterning
MinD
LifeAct
lag (s)
0
10
kymo.
30s
+rap
intensity
50s
6s lag
corr.
t=0
30s
D
image level timeseries
image level x-correlation
pixel-level examples from image
dynamic actin spots
dynamic region
MinD
LifeAct
lag (s)
0
20
spots kymo.
30s
+rap
intensity
30s
10s lag
corr.
t=0
30s

**Figure S11. Additional examples and analysis of MinD-ActA driven actin patterning using dynamic MinDE control signals.**

(A) Additional examples of MinDE to ActA patterning circuits in which the MinDE control signal is dynamic, with waves of MinD driving oscillatory ActA recruitment. These control signals are inducibly connected to the ActA effector protein upon addition of rapamycin, which will stimulate actin polymerization through recruitment of Arp2/3 and VASP. See also Movie S22.

(B) Example of a cell with a smooth MinDE wave control signal. Representative image showing MinD and LifeAct distribution in the cell. A pixel-level raw time-series data for the two signals shows a clear time-lag between the two signals, as does a kymograph of the FIR-filtered versions of the LifeAct and MinD signals. A representative cross-correlation profile for two pixel-level signals is shown, with the extracted time-lag (argmax=6s) indicated. This results in an image-level cross-correlation profile that shows a consistent 6s time-lag everywhere within the cell. See also Movie S23.

(C) As in (B), but for a cell with a more turbulent MinDE control signal. Representative image showing MinD and LifeAct distribution in the cell. A pixel-level raw time-series data for the two signals shows a clear time-lag between the two signals, as does a kymograph of the FIR-filtered versions of the LifeAct and MinD signals. A representative cross-correlation profile for two pixel-level signals is shown, with the extracted time-lag (argmax=6s) indicated. This results in an image-level cross-correlation profile that shows a consistent 6s time-lag, but its overall distribution within the cell is patchier. See also Movie S23.

(D) As in (B) but for a cell that contains a mixture of both static and dynamic MinDE control signals (contrast to the static spots analysis of the same cell in Fig. S9B). Representative image showing MinD and LifeAct distribution in the cell. A pixel-level raw time-series data for the two signals shows a clear time-lag between the two signals, as does a kymograph of the FIR-filtered versions of the LifeAct and MinD signals. A representative cross-correlation profile for two pixel-level signals is shown, with the extracted time-lag (argmax=10s) indicated. This results in an image-level cross-correlation profile that shows a consistent 10s time-lag only in the regions of the MinDE control signal that are dynamic. See also Movie S23.

**Supplemental Movie Legends.**

A playlist containing all supplementary movies in a streaming format can be found at: http://youtu.be/watch?v=KDgHVrjdydY&list=PLx2v1NUlEZF_idH1goDHYimtlP8c7mnLB&index=1

**Movie 1. The MinDE reaction-diffusion system generates diverse protein dynamics and patterns across a range of mammalian cell lines.** Example time lapse imaging of reaction-diffusion protein dynamics and patterns seen in populations of 3T3, U2-OS, 293T, and K562 cells expressing a range of levels of mCherry-MinD and MinE-GFP.

**Movie 2. Additional detail for different classes of MinDE protein patterning behavior in mammalian cells.** Composite time-lapse fluorescence images of MinDE reaction-diffusion phenomena, including waves, standing oscillations, turbulent patterns, and spotted domains.

**Movie 3. Synthetic spatiotemporal protein patterns in mammalian cells using the MinDE reaction-diffusion system: traveling waves.** Example composite images, kymographs, and pixel-level quantification of traveling waves.

**Movie 4. Synthetic spatiotemporal protein patterns in mammalian cells using the MinDE reaction-diffusion system: standing oscillations.** Example composite images, kymographs, and pixel-level quantification of standing waves.

**Movie 5. Synthetic spatiotemporal protein patterns in mammalian cells using the MinDE reaction-diffusion system: persistent "leopard print" patterns.** Example composite images, kymographs, and histogram of inter-spot distances.

**Movie 6. Examples of other unique MinDE protein patterns in mammalian cells.** Example composite images, kymographs, and pixel-level quantification of fast turbulent waves and slow gliding waves. TIRF microscopy shows MinDE circuit activity at adhered regions of the cell.

**Movie 7. MinDE patterning is supported by the endomembrane system.** MinDE signals colocalize with the endomembrane. Hilbert transform derived oscillatory power representations are consistent with MinDE activity being supported endomembrane system.

**Movie 8. Instantaneous signals generated by Hilbert transform distinguish traveling waves and standing oscillations.** Hilbert transform derived instantaneous phase and power representations distinguish dynamic MinDE patterns. Traveling waves have smooth phase field gradients and uniform power while standing oscillations display alternating nodal structures and do not have oscillatory power in regions between nodes.

**Movie 9. MinDE phase field reports on spiral centers where there is no MinDE oscillation.** Hilbert transform derived instantaneous phase fields are used to calculate vorticity. Regions where vorticity is approximately $2\pi$ or $-2\pi$ are spiral centers. The positivity or negativity of the vorticity value indicates the directionality of the rotation. The spiral centers are displayed on the phase field and are color-coded to show clockwise (pink), and counterclockwise (blue) spiral centers located with the cell.

**Movie 10. MinDE circuits are "cellular radios" that broadcast FM-barcoded single cell fluorescence data.** Single fluorescence channel MinDE images encode oscillatory time domain signals that can be transformed to frequency domain signals using fast Fourier transform. Tuning to frequencies within an image enables spectral separation of cells with MinDE circuits operating at that frequency. Images false colored by frequency enable visualization of spectral separation of overlapping cells. FM-barcoding provides a unique signal to track individual cells across time in densely populated cellular environments.

**Movie 11. FM-barcoded 3T3 cells tracked over 24 hours.** Widefield 3T3 fluorescence time lapse images recolored by MinDE frequency. Images were collected for 3 minutes at 1 FPS every 30 mins for 24 hours.

**Movie 12. MinDE circuits frequency barcode cell identity across a broad range of cell types.** MinDE circuits are active when installed in 3T3, U-2 OS, 293T, and K-562 cells. Frequency domain image processing can isolate, segment, and FM-barcode cells with active MinDE circuits regardless of cell type or shape.

**Movie 13. Digital signal processing enables separation of different oscillations in different compartments of the same cell.** Example of spectral separation of MinDE signals occurring in different sub-compartments of the same cell.

**Movie 14. Hilbert transform analysis pipeline extracts instantaneous MinDE signal content.** HT generates an analytic MinDE time signal that reports on instantaneous MinDE signal content. The angle of the analytic signal generates the instantaneous phase representation, and the magnitude of the analytic signal generates the instantaneous amplitude representation. The phase imparts information on the momentary structure of the MinDE pattern in an instant in time while the amplitude imparts information on the oscillatory power. The instantaneous frequency is generated by taking the derivative of the unwrapped phase time signals.

**Movie 15. Strategic use of FFT, FIR filters, and HT enables separation of subcellular compartments with unique MinDE signals.** The pixel-level power spectrum of mCherry-MinD signals collected over a 6-hour time series revealed distinct frequencies corresponding to separate nuclear and cytoplasmic MinDE oscillations. Using Finite Impulse Response (FIR) filters based on the power spectrum enable separation of the nuclear and cytoplasmic signals. Hilbert Transform was used to estimate the instantaneous power of each signal throughout the 6-hour recording. This produces a multi-channel representation of the data that separately labels and tracks the cytoplasmic and nuclear signals.

**Movie 16. Continuous wavelet transform is complementary to Hilbert transform analysis.** Continuous wavelet transform yields instantaneous phase, frequency, and power information for MinDE signal content identical to Hilbert transform.

**Movie 17. CWT separates subcellular MinDE signals.** CWT enables separation and visualization of subcellular structures. However, the CWT does not require pre-processing to achieve this. CWT separates cytoplasmic and nuclear signals into different scale frequencies across the entire duration of the time series data. By visualizing the different scale frequencies that contain power for signals of interest, it is possible to directly isolate cytoplasmic and nuclear signal content.

**Movie 18. Proteins payloads can be readily coupled to MinDE signals.** A plug-and-play platform for inducible recruitment of protein payloads to MinDE circuits based on rapamycin-dependent interaction between FRB-mCherry-MinD and FKBP-payloads. Inducible recruitment of a simple BFP payload is shown.

**Movie 19. Protein payloads can dynamically modulate MinDE reaction-diffusion behavior in real time.** Example composite images and kymographs of protein payloads modulating MinDE reaction-diffusion behavior. RGG, FUS, and DDX4 are LLPS modules with varied multivalent macromolecular interaction strength (RGG < FUSN < DDX4). Loading the LLPS module onto the MinDE circuit causes real-time frequency modulation of reaction-diffusion behavior where decrease in frequency is proportional to the strength of the LLPS modules. TPPP binds microtubules. Loading TPPP onto the MinDE circuit sequesters MinD to the cytoskeletal microtubules and abolishes reaction-diffusion behaviors.

**Movie 20. Broadcasting PKA activity dynamics on MinDE reaction-diffusion carrier signals.** MinDE-PKA signaling reporter circuit shows changes in PKA activity dynamics through coupling of mCerulean-FHA1 fluorescence signals to MinD oscillatory signals.

**Movie 21. MinDE signals broadcast single cell frequency-barcoded signals that report host-cell PKA activity dynamics.** PKA signaling trajectories were computed at a single-pixel level and aggregated at the cellular level based on FM-barcode using the frequency-domain cell segmentation strategy.

**Movie 22. ActA recruitment to MinDE circuits drives actin polymerization.** Example composite images that show MinDE signals can direct ActA organization to promote synthetic actin polymerization.

**Movie 23. MinD-ActA driven actin patterning using dynamic MinDE control signals.** FRB-FKBP interaction couples ActA activity to MinDE signals. LifeAct-GFP reports on subcellular actin polymerization. In addition to static actin patterns, MinDE signals can also direct dynamic waves of actin polymerization. A pixel-level raw time-series data for the two signals shows a clear time-lag between the two signals. A representative cross-correlation profile for two pixel-level signals is shown, with the extracted time-lag (argmax=8s) indicated. Image-level cross-correlation profile that shows a consistent 8s time-lag only in the regions of the MinDE control signal that are dynamic.
